# On the mechanical origins of waving, coiling and skewing in *Arabidopsis thaliana* roots

**DOI:** 10.1101/2023.05.17.541081

**Authors:** Amir Porat, Arman Tekinalp, Yashraj Bhosale, Mattia Gazzola, Yasmine Meroz

## Abstract

By masterfully balancing directed growth and passive mechanics, plant roots are remarkably capable of navigating complex heterogeneous environments to find resources. Here we present a theoretical and numerical framework which allows us to interrogate and simulate the mechanical impact of solid interfaces on the growth pattern of plant organs. We focus on the well-known waving, coiling and skewing patterns exhibited by roots of *Arabidopsis thaliana* when grown on inclined surfaces, serving as a minimal model of the intricate interplay with solid substrates. By modelling growing slender organs as Cosserat rods that mechanically interact with the environment, our simulations verify hypotheses of waving and coiling arising from the combination of active gravitropism and passive root-plane responses. Skewing is instead related to intrinsic twist due to cell file rotation. Numerical investigations are outfitted with an analytical framework that consistently relates transitions between straight, waving, coiling and skewing patterns with substrate tilt angle. Simulations are found to corroborate theory and recapitulate a host of reported experimental observations, thus providing a systematic approach for studying *in silico* plant organs behavior in relation to their environment.

**Significance:** Plant roots exhibit an exceptional ability to navigate in heterogeneous soil environments while overcoming obstacles. Our study combines theory and experimental observations to interrogate and simulate the mechanical impact of obstacles on organ growth. As a test case we focus on well-known observations of waving, coiling and skewing growth patterns of *Arabidopsis thaliana* roots grown on inclined substrates. Overall, our study explains a broad set of experimental observations through the minimal ingredients of gravitropism and passive mechanics. Our numerical framework provides an *in silico* laboratory, yielding quantitative insight into the dynamics of growing organs at the intersection of active processes and passive mechanics, applicable beyond plants to any slender growing system, from neurons or fungal hyphae to novel soft robots.

---

The ability of roots to grow in soil is critical for plant health and crop yield, enabling the uptake of water and nutrients, providing anchorage and stability of eroding soil (1–4). This is no small feat. Indeed, soil is a highly heterogeneous environment, characterized by non-uniform concentrations of resources as well as obstacles such as rocks and compacted soil. In negotiating with the environment, plant roots combine passive physics and active growth-driven mechanisms, termed tropisms, whereby dedicated organs sense stimuli such as water (hydrotropism) or gravity (gravitropism), and redirect growth appropriately. While dynamics of tropisms are generally understood (5, 6), and aspects of root mechanics and their effect on growth have been described (7– 11), a consistent framework to dissect the interplay between passive and active responses, heterogeneous environments and ensuing growth patterns, is still missing (12–14). The essence of this complex interaction is on display in controlled experiments of roots growing on inclined agar gel substrates, whereby remarkably different behaviors are observed to emerge: roots grow straight on vertical planes, grow in waving patterns as tilt increases, skew in some cases, and eventually coil when tilt approaches the horizontal plane (Fig. a). These growth patterns are well-documented in genetically driven phenotypes of *Arabidopsis thaliana* (15–21). Although insightful, such genetic approaches are limited to given species, and do not formally address the fundamental role of root and environmental mechanics. Thus, while gravitropic responses and root-substrate mechanics clearly play a role (17, 22), underlying mechanisms remain subject of debate. For instance, both circumnutation (16, 23– 25), the intrinsic circular movements of root tips, and thigmotropism (15, 22, 26), the active response to touch, have been suggested to be at the basis of waving and skewing patterns. Here, we combine advances in the modeling of growing rod-like organs (25, 27–36) and 3D numerical simulations (37) to gain broader insight.

## Results

### Modeling of root mechanics and growth

In modeling the waving and coiling experiments of *Arabidopsis thaliana*, we begin by assuming separation of timescales between slow growth-driven root responses and fast mechanical relaxation (Fig. d), allowing us to decouple the two processes in a quasistatic manner (28, 29).

Slender roots are represented as Cosserat rods (37), which are 1D elastic continuous elements able to undergo all modes of deformations – bending, twisting, stretching, shearing – and reconfigure in 3D space (Fig. b). We mathematically describe a slender rod by a centerline **r**(*s, t*) ∈ ℝ^3^ and a rotation matrix **Q**(*s, t*) = {**d**_1_, **d**_2_, **d**_3_}^−1^, providing a local material frame. When the rod is at rest, its length is *L* and the corresponding material coordinate is *S* ∈ [0, *L*], while *l* and *s* ∈ [0, *l*] denote the length and arc-length of the deformed (stretched) filament, and *t* is time. If the rod is unsheared, **d**_3_ points along the centerline tangent 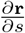, while **d**_1_ and **d**_2_ span the normal–binormal plane, i.e. the cross-section. Shearing and stretching shift **d**_3_ away from 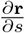, quantified by the shear vector 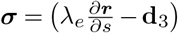, where *λ*_*e*_ ≡ ∂*s/*∂*S* is the local elastic stretch. The curvature vector ***κ*** = *κ*_1_**d**_1_ + *κ*_2_**d**_2_ + *κ*_3_**d**_3_ encodes the rotation rate of the local frame along the material coordinate so that 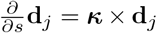. We define the bending stiffness matrix **B** and shearing stiffness matrix **S** in the rest configuration. Then, the mechanical equilibrium of a rod-like root is described by

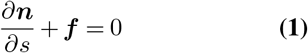

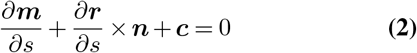

where ***n*** = **S**(***σ*** − ***σ***^0^) represents internal forces, related to shear and elastic stretch of the centerline, ***m*** = **B (*κ*** − ***κ***^0^)represents internal torques, related to bending and twisting, and ***σ***^0^ and ***κ***^0^ are intrinsic strains. As detailed in (37), incompressibility is incorporated by rescaling **B, S** and ***κ*** through *λ*_*e*_ such that 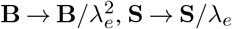 and ***κ*** → ***κ****/λ*_*e*_. Lastly, external forces ***f*** and torques ***c*** (per unit length) capture overall environmental effects.

Next, we outfit this model with active gravitropic dynamics driven by differential growth (30, 38, 39). Axial growth is implemented via the additional stretch factor *λ*_*g*_, defined with respect to an initial material coordinate *S*_0_ ∈ [0, *L*_0_], such that *L*_0_ is the initial length and *λ*_*g*_ ≡ ∂*S/*∂*S*_0_ (Fig. d). By denoting time derivatives as 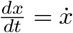, we introduce the average relative growth rate as 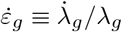. Root growth is restricted to a finite growth zone of length *L*_gz_ extending from the tip (Fig. e), reflecting experimental evidence (8, 40, 41).

Tropic movements are directed by environmental cues, which are generally perceived via dedicated sensory systems. Gravity in particular is sensed via specialized cells close to the tip (42), and translated into a redistribution of growth hormones across the root cross-section. This results in axial differential growth, which in turn leads to a change in the root’s curvature, redirecting the organ towards the stimulus. This machinery is mathematically captured via the local differential growth vector **Δ**(*s, t*) (30, 38, 39)

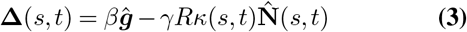

where the first term represents the gravity stimulus, with ***ĝ*** the direction of gravity and *β* the gravitropic gain or sensitivity (38). The second term represents the *prioprioceptive* response. This can be thought of as a counter-curving response that balances the gravitropic one, based on the sensing of the organ’s own shape (38). Here *γ* is the proprioceptive gain, 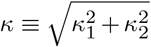 is the norm of the bending curvature, and 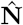 is the normal director of the center-line’s Frenet-Serret frame (Fig. c). Given the average growth rate 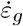 and the organ radius *R*, we connect the change in local intrinsic curvature ***κ***^0^(*s, t*) within *L*_gz_, to the differential growth **Δ**(*s, t*) projected on the cross section (30, 38, 39)

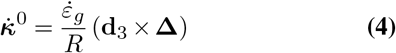

as detailed in SI Appendix 2. For simplicity, here we drop the explicit dependence on (*s, t*). Thus, Eqs. 3 and 4 capture gravitropic dynamics, relating sensing (*β****ĝ***) to differential growth (**Δ**), which in turn feeds into the root shape (***κ***^0^). As described in the Methods section, **Δ** can be rewritten to include different types of internal and external cues, such as phototropism or circumnutations, as well as different sensing profiles (apical and local). Finally, to distill the essential mechanisms underlying the range of observed growth patterns, we neglect the effect of substrate deformations (17, 43), and the coupling between root growth, static friction, and stick-slip dynamics (41, 44, 45). These elements nonetheless deserve future attention.

Some *Arabidopsis* mutants (46–48) exhibit a cell profile which twists during growth (Fig. b). We model this additional effect by allowing the material frame in the growth zone *L*_gz_ to twist around the centerline of the organ. This twist is described by an increasing register angle *ξ*, characterized by the angular velocity at the apex

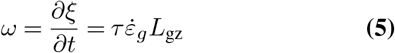

where *τ* is a constant intrinsic twist line density (Fig. c).

Finally, contact forces that prevent the root from penetrating the substrate, and adhesion forces that maintain the mature region fixed in place (mimicking the effect of root hair attachments and lignification, Fig. a) are encapsulated in **f**, as detailed in Methods and SI.

Overall, Eqs. 1-5 define our model, which we numerically discretize and solve using the open-source software *Elastica* (49), demonstrated across a range of biophysical problems (50–55) entailing fiber-like structures (Methods and SI Appendix 3).

### Simulations recapitulate waving and coiling patterns

Based on waving and coiling experiments (15–21), we simulate *Arabidopsis thaliana* roots growing on a tilted plane for a range of angles *α* (from vertical to horizontal), as well as gravitropic and proprioceptive gains *β* and *γ*. We use parameters and mechanical properties typical of *Arabidopsis* (Table 1 in Methods). For simplicity we assume frictionless growth while fixing the sessile (mature) zone to the substrate (17, 43), as discussed above. At this stage, we neglect root twisting during growth, although we will revisit it in later sections.

**Table 1.** Parameters used to emulate *Arabidopsis thaliana* roots. The values taken have the same order of magnitude as their measured values, as described in (8, 27, 40, 58, 67). The values for *Rτ* are chosen based on data from (23, 26, 60–62) and Eq. 27 as plotted in SI Appendix 9. The last relation was used in order to fit experimental data in Fig..

| Parameter | Value used | Units |
| --- | --- | --- |
| $R$ | 100 | $\mu\text{m}$ |
| $L_{gz}$ | 1 | mm |
| $\dot{\epsilon}_g^0$ | 0.2 | 1/h |
| $E$ | 50 | MPa |
| $\beta$ | 0.05-1.0 | |
| $\gamma$ | 0.0-10.0 | |
| $R\tau$ | 0.0-0.8 | |
| $v_g^{\text{tip}} = \dot{\epsilon}_g^0 L_{gz}$ | 0.2 | mm/h |
| $\beta L_{gz}/R$ | 0.5 | |

As illustrated in Fig. a and supplementary movies 1, 2, simulations capture observed root behavior. Indeed, simulated roots grow straight on a vertical plane (*α* = 0), and transition to waving as the substrate tilt angle reaches *α* ≈ 15^0^, consistent with experimental measurements (Fig. a). A further increase in tilt angle initiates a second phase transition, around *α* ≈ 50^0^, and roots begin to coil, again consistent with experiments (Fig. a). Transition angles are also consistent with experimental values (43) (Fig. e), as we shall elaborate later.

To gain intuition about the waving and coiling process, as well as their transitions, we analyze the system from an energetic and topological perspective. Tip dynamics are tracked via the planar angle *θ*^tip^. This is the angle between the projections of gravity and the tangent of the apex on the tilted plane (Fig. a and c). The evolution of the stored energy in the growth zone is described through the approximate elastic bending energy 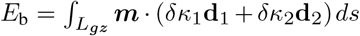 and elastic twist energy 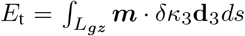, where *δ****κ*** = ***κ****/λ*_*e*_ − ***κ***^0^ and 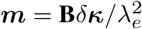 (Fig. d). Additionally, in order to concisely capture root morphology and its reconfigurations, we consider the topological quantities writhe (Wr) and twist (Tw) in the growth zone (Fig. e). Writhe is computed by treating the growth zone as an open loop (56), and is used here as a measure of out-of-plane bending of the growth zone’s centerline. If Wr = 0, then the centerline is in-plane, while positive or negative values of Wr correspond approximately to right- or left-handed out-of-plane bending, respectively (SI Appendix, Fig. S2). Twist instead determines the rotation of the local frame around its centerline. Positive and negative values correspond to right-handed and left-handed rotations accordingly (SI Appendix, Fig. S2). In a closed or infinitely long rod, the geometric integral quantities Wr and Tw are related topologically, and their sum is constant and constrained by the number of formed loops (57). In our case their sum is not conserved, however their topological connection is useful to qualitatively understand the system dynamics.

As can be seen in Fig. a, the onset of waving and coiling follows a first turning event. Fig. b shows snapshots at different stages of a waving organ, animated in supplementary movie 1 and marked for reference in Figs c-e. Snapshot (i) occurs just before the first turning event, where the root grows in a plane defined by its initial tangent and the direction of gravity, as represented by the constant values of *θ*^tip^ (Fig. c), Wr and Tw (Fig. e). Due to gravitropism, the root tends to grow into the substrate (Fig. d), which in turn resists to growth, leading to an increase in bending energy *E*_*b*_ (Fig. d). This energy is eventually released, breaking symmetry, in the form of an elastic instability - snapshot (ii) - whereby the root tip slips or tilts to one side, bending and twisting. This process is reflected by a sharp drop in *E*_*b*_ accompanied by a jump in all other parameters (Fig. c-e). We note that while in simulations this initial symmetry breaking is purely caused by an elastic instability, in experiments a number of factors may further contribute to it, from geometrical or material imperfections to small intrinsic root twists.

We continue to follow the growth of the root during a subsequent turning event. Snapshot (iii) occurs right after the second turn where the tip angle *θ*^tip^ is at a maximum, and the bending energy *E*_*b*_ is at a minimum. For further insight, it is useful to decompose **Δ** into components parallel and perpendicular to the substrate. The parallel component causes the root to reorient along the projection of gravity on the plane, leading to a decrease in the planar tip angle *θ*^tip^. The perpendicular component pushes the apex into the plane, leading again to an increase in *E*_*b*_, similar to the initial symmetry breaking and in line with observations of substrate deformations (17). As new cells are produced at the tip, older cells stop growing and become part of the mature zone. Thus, accumulated twist leaves the growth zone, so that, at snapshot (iv), *E*_*t*_ reaches a minimum while *E*_*b*_ reaches a maximal value. Then, as for the first bending event, the organ releases bending energy by twisting, and *E*_*b*_ is converted into *E*_*t*_. This elastic relaxation is accompanied by a conversion of Wr into Tw, since these quantities are topologically related (56). Both *E*_*t*_ and Tw reach a maximum value at snapshot (v). At this point Tw is convected out of the growth zone faster than it is generated, and Tw and *E*_*t*_ decrease. The organ is now back to stage (iii), and the process repeats itself. Since the gravitropic correction of curvature is slower than the elastic relaxation, *θ*^tip^ repeatedly overshoots in the direction of gravity, producing the observed waving patterns.

We note that our simulations show that neither thigmotropism nor circumnutation are strictly required in order to produce waving and coiling, unlike previously argued (15, 16, 23, 24, 26).

#### Scaling analysis

We now perform a scaling analysis of the growth dynamics described in Eqs. 3 and 4, and relate model parameters to experimentally measurable variables: wavelength *λ* and amplitude *A* of root waving patterns, as defined in Fig. c. Comparing predicted values to experimental measurements will allow to quantitatively corroborate our model. In order to derive relations for amplitude and wavelength, we focus on the curvature of the waving pattern. Simulations (Fig.) show that the elastic stresses associated with bending, due to the organ pushing into the substrate, are accumulated until they relax by twisting sideways, which rotates the curvature vector. This suggests that the maximal value of the in-plane curvature of the waving pattern *κ*, obtained right after twisting, may be comparable to the maximal curvature 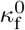 induced by the gravitropic response in a free organ tilted at the same angle *α* of the substrate. Thus, we hypothesize that

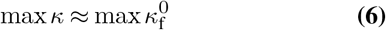

In SI Appendix 6 we provide a detailed analysis supporting this assumption, decomposing the dynamics of a tilted organ into components parallel and perpendicular to the tilted plane. This assumption is also supported by Fig. c, which compares the oscillating tip angle of a root waving on a substrate (blue) with the gravitropic bending of a corresponding free organ (pink). This comparison further suggests that the wavelength *λ* of the waving pattern is dictated by the damped oscillations inherent to the free organ’s gravitropic dynamics (58), while the amplitude *A* is modulated by the mechanical interaction with the substrate.

Therefore, in order to find *λ*, we first evaluate the characteristic gravitropic turning period *T*_0_, corresponding to the time a tilted free root takes to turn towards the direction of gravity (27, 38). We rewrite Eq. 4 in non-dimensional form, assuming *γ* = 0 for simplicity, and rescaling length and curvature by *L*_gz_, yielding 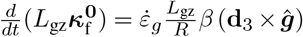. Values in parentheses are dimensionless, as well as the coefficient 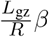. Since **d**_3_ and ***ĝ*** are unit vectors, we expect the turning time *T*_0_, related to the rate of change of the normalized curvature 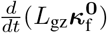, to scale as a function of 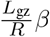, with 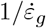 the overall timescale of the problem. Based on free organ’s simulations (see SI Appendix 5) we find

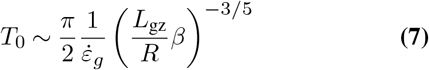

Here *L*_gz_*/R* is the geometric slenderness ratio of the growth zone, and we interpret 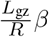 as an effective gravitropic sensitivity (58). We approximate 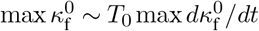 and substitute 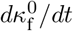 from Eq. 4 (with *γ* = 0) to obtain

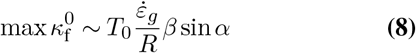

This equation provides an estimate for 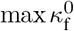, which together with Eq. 6 relates the waving curvature to the gravitropic curvature. In Fig. a we validate Eq. 6 for a range of tilt angles by plotting max *κ*, the maximal curvature measured from simulated waving patterns, against 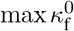, the analytically predicted maximal curvature for a free tilted organ in Eq. 8, finding good quantitative agreement.

Having verified that the timescale of the waving pattern is related to the gravitropic bending, we can approximate the length *λ/*2 grown during a turning event (Fig. c) as

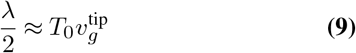

where 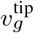 is the tip growth speed. Then, following simple geometric arguments, we relate the amplitude *A* to the wavelength. After a turn event, 2*A* and *λ/*2 form an orthogonal triangle (Fig. c). Based on Eq. 6, and since in the gravitropic response of a free organ the angle between the base and the tip is equal to the tilt angle *α* (Fig. d), we can infer

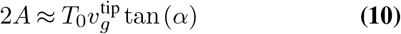

By combining Eq. 9, 10, we then obtain

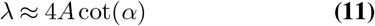

directly relating wavelength *λ* and amplitude *A* through the incline *α*. We compare Eq. 11 with simulations and experiments (16, 43, 59) in Figs. b, noting consistent trends. The individual scaling relations (Eqs. 9,10) are also tested in SI Appendix 5. Throughout, we assume local sensing. We additionally report results for apical sensing in SI Appendix 5 (see Methods), observing no qualitative differences.

#### Transitions between straight, waving and coiling

Here we provide an intuitive rationale for the two critical tilt angles at which the root transitions from straight to waving (*α*_s→w_) and from waving to coiling (*α*_w→c_), with the full derivation reported in SI Appendix 7.

Simulations of Fig. suggest that waving patterns are initiated by an elastic instability, which occurs when it becomes energetically convenient to twist rather than to bend, i.e. *E*_t_ *< E*_b_. Again assuming that the intrinsic curvature is captured by the gravitropic response of the free organ max 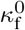 (Eq. 8), before the turn we can approximate 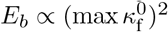. The accumulated angle during the gravitropic response is roughly equal to the tilt angle 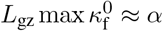, meaning that *E*_*b*_ increases with tilt angle. However *E*_t_ is constant, and at a critical angle *α*_s→w_ the two energies intersect *E*_t_ = *E*_b_(*α*_s→w_) and waving occurs. The critical angle (detailed in SI Appendix 7) follows

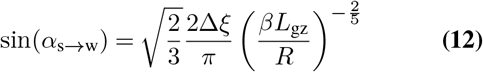

where Δ*ξ* is a fitting parameter representing the change in the register angle *ξ* created by the integrated elastic twist (Fig. c). The transition between waving and coiling is instead related to the ratio between (i) the time to turn towards gravity after a twist induced relaxation and (ii) passive orientation drift (27) – the rate at which the tip angle of a curved organ increases due to growth, maintaining the same curvature, thus reorienting growth uphill in this case. The former is proportional to the amplitude of the waving pattern *T*_0_ tan (*α*) in Eq. 10, and the latter follows 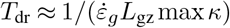 (27). At at a critical angle *α*_w→c_ orientation drift occurs faster than turning, *T*_dr_ *< CT*_0_ tan (*α*), and coiling ensues. The coiling critical angle (detailed in SI Appendix 7) then follows

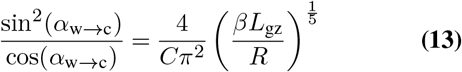

where *C* is a fitting parameter.

We corroborate the predicted critical angles both numerically and experimentally. Fig. d shows the *configuration space* of simulated organs over a range of *α* and effective gravitropic bending *βL*_gz_*/R* (where for *Arabidopsis* roots we use 0.5, see Table 1). For each set of parameters we compare the conformation observed in simulations (◼ symbols for straight, ♦ for waving,• for coiling) to the predicted one (represented by background colors, separated by the critical angles), with Δ*ξ* = *π/*8 and *C* = 0.5, finding good agreement. Configuration spaces for cases with *γ* = 1, and apical sensing, can be found in SI Appendix 7, where we note no qualitative difference. In Fig. e we also compare our results to the coiling probability measured *in vivo* by Zhang et al. (43), again finding good quantitative agreement.

#### Intrinsic twist yields skewing patterns

Lastly, we focus on the skewed waving patterns observed in some *Arabidopsis* mutants (26, 60–62) (Fig. a). Observations suggest that the skewing angle is related to the twisting of root cell files (e.g. Fig. b), which may be represented as an intrinsic twist of the material frame around the centerline of the growth zone, as described in Eq. 5 and detailed in the Methods (Fig. c). When accounting for such rotations (*ω* in Eq. 5), simulations recover observations of skewing (Fig. a and supplementary movies 3-5). We see how increasing the ratio between the bending time and the twisting velocity, *ωT*_0_, results in larger skewing angles and washes out the waving pattern. This is reflected in the evolution of energetic and topological variables: *ωT*_0_ = 1 (Fig. d-f) exhibits oscillatory behavior similar to that of regular waving patterns, while *ωT*_0_ = 8.3 (SI Appendix, Fig. S10) transitions to a more monotonic behavior. Here, the initial breaking of symmetry manifests as a smooth, twist-induced rotation, facilitated by the intrinsic cell file rotation (26). We also note that circumnutations are not required, however they could have a secondary role, as we find that with no intrinsic twist circumnutations lead to disordered patterns and low skewing angles (SM 8), explaining observed coiling in agravitropic roots (16).

Next, we relate the average skewing angle ⟨*θ*⟩ (defined in Fig. a) to *ω*, for two limiting cases. When *ωT*_0_ ≪ 1, the apical rotation due to intrinsic twist acts as a small perturbation to the waving mechanism, accelerating turns in the same direction, while slowing down opposite turns (Fig. d-f). Thus, the amplitude of the turn increases in one direction, *A*_+_ ≈ *A*(1 + *ωT*_0_), and decreases in the other, *A*_−_ ≈ *A*(1 −*ωT*_0_). From trigonometric arguments sin⟨*θ*⟩ is the horizontal displacement due to the difference in amplitude over two turns, divided by the wavelength sin⟨*θ*⟩ ≈ Δ*A/λ*, with Δ*A* = 2*A*_+_ − 2*A*_−_ = 4*AωT*_0_. Substituting the expressions for *λ* and A in Eqs. 9-10 yields

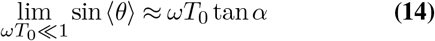

In the opposite limit *ω*_0_*T*_0_ ≫ 1, the twist-induced rotation dominates, and the waving pattern disappears altogether. Following simple geometric arguments (Fig. d), the skewing angle ⟨*θ*⟩ reduces the in-plane gravitropic component *β*_∥_ = *β* cos *α* sin ⟨*θ*⟩, however it does not affect the perpendicular component directed into the plane, *β*_⊥_ = *β* sin *α*, which causes the apex to drift uphill when twisted into the plane. The constant skewing angle suggests that these two competing processes balance each other, *β*_⊥_ = *β*_∥_, yielding

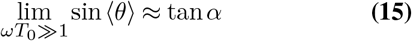

We corroborate our skewing model (limiting relations Eqs. 14, 15 and simulations) against experimental observations (26, 60–62) in Fig. g. As can be seen, theory and simulations are found to be consistent among themselves as well as able to capture general trends across a variety of Arabidopsis mutants. Only a particular strain, expressing MBD-GFP (26), is found to significantly depart from our predictions (top/left corner of Fig. g). While we cannot comment on the effect of this specific mutation on growth behaviour, we note that these outliers are themselves a subset of the specimens investigated in (26), which are otherwise in agreement with all data of Fig. g.

## Conclusions

We developed a numerical framework for simulating the interaction of plant roots with solid interfaces, integrating passive mechanics and active growth-driven mechanics. Our methods are shown to reproduce the range of growth patterns exhibited by *Arabidopsis* roots grown on a tilted plane, a well documented biophysical plant model. Our simulations illustrate how the interplay between active gravitropic response and passive elasticity are the minimal requirements to generate straight, waving and coiling patterns, while the addition of intrinsic twist is responsible for skewing. We find that neither thigmotropism nor circumnutation are required to recover waving or skewing patterns, as previously argued (16, 23– 25), however they could play a secondary role.

A scaling analysis reveals that the amplitude of waving patterns is modulated by the mechanical interaction with the substrate, while the wavelength depends on oscillations inherent to the gravitropic dynamics of a free organ, regardless of the interaction with the plane. Based on this analysis we develop analytical expressions for the critical tilt angles for which the root behavior transitions from straight to waving and then coiling. Further, we elucidate the relation between skewing angles and intrinsic twist. We corroborate all these analytical insights by comparing model predictions to both simulations and experimental observations, obtaining good agreement.

This framework provides, effectively, an *in silico* laboratory to form and test hypotheses relative to the behaviors of plants and their mechano-sensory machinery in realistic environments. Future extensions will consider heterogeneous terrain via the inclusion of granular mechanics. Finally, we note that since our approach is general and agnostic to the underlying biological building blocks, it can be applied to growth-driven systems other than plant organs, such as neurons, fungal hyphae, or new generation of growing robots (63–66).

## Materials and methods

Here we consider a plant organ as a slender rod, whose centerline is parameterized by *s*, with *s* = 0 at the base and *s* = *l* at the tip, *l* being the organ length (Fig. b). In order to describe its dynamics, we assume a separation of timescales between slow growth-driven processes and fast elastic relaxation (28, 29). At each time step of our integration scheme we first update the configuration of the organ according to the growth process alone, and then allow it to relax mechanically following Cosserat rod theory (37) (Fig. d). For the sake of clarity, the intermediate stress-free configuration is parameterized with *S*, in order to differentiate it from the final relaxed configuration described with *s*. The initial configuration is marked with *S*_0_. In what follows we bring the governing equations describing the growth dynamics and the mechanical relaxation.

### Shape

At each point *s* along the centerline we define the position vector ***r***(*s, t*), which provides the 3D description of the rod at time *t* (Fig. b). We also define a local orthonormal material frame {**d**_1_(*s, t*), **d**_2_(*s, t*), **d**_3_(*s, t*)}, where **d**_1_ and **d**_2_ span the cross-section of the organ, and in a shearless and inextensible system **d**_3_ coincides with the tangent of the centerline. The local curvature vector ***κ***(*s, t*) (or Darboux vector) of the center-line is defined through the the relation 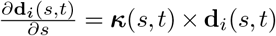 (37). The components of curvature projected along the principal vectors of the material frame (***κ*** = *κ*_1_**d**_1_ + *κ*_2_**d**_2_ + *κ*_3_**d**_3_) coincide with bending (*κ*_1_, *κ*_2_) and twist (*κ*_3_) strains in the material frame. Introducing elastic stretch may lead to a difference between the arc-length configuration *s* and the stress-free configuration *S*, described by the local stretch (37) 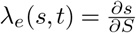. Shear may leads to an incongruity between **d**_3_ and the tangent of the center-line, and the local deviation is described by the translation vector 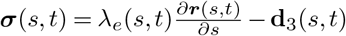 (Fig. b).

### Growth

Root growth occurs in the *growth zone*, an area of length *L*_gz_ just below the root tip. Cells divide at the tip, and elongate within the growth zone, until stop elongating and reach the *mature zone* (Fig. e). The initial configuration then represents the material (or Lagrangian) coordinate which flows due to growth with velocity *v*_*g*_, which is the integral of the mean axial relative growth rate 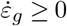

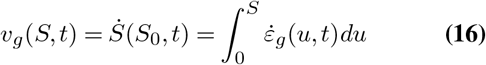

Here the growth rate is the time derivative of the logarithmic growth strain *ε*_*g*_ = ln *λ*_*g*_, where 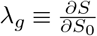 is the growth stretch. The increase in rest lengths enters the dynamics via the actual and intrinsic stretch vectors, defined as 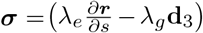 and ***σ***^0^ = −(*λ*_*g*_ − 1) **d**_3_ respectively, such that 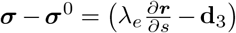. As described in the main text, growth-driven movements are generally classified into tropic and nastic movements, where the former are due to external stimuli, such as gravitropism, and the latter are due to internal cues, such as the oscillatory movements of circumnutations. Changes in curvature are due to the anisotropic redistribution of growth hormones, leading to asymmetric growth along the cross-section captured by the differential growth vector **Δ**(*s, t*) (30). The dynamics resulting change in intrinsic curvature ***κ***^0^ for *L*(*t*) − *L*_*gz*_ ≤ *s* ≤ *L*(*t*) within the growth zone follows (30)

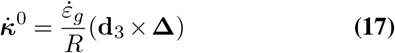

Here 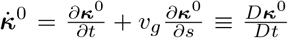 is a material derivative which accounts for the growth of the centerline. For simplicity we assume no intrinsic twist, 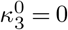. The cross product **d**_3_ × **Δ** represents a projection of **Δ** on the local cross-section of the organ (details in SI Appendix 2). The tropic and nastic movements then contribute to the total differential growth vector **Δ**. In the case of gravitropism we have

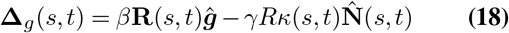

The first term represents the gravity stimulus, with ***ĝ*** the direction of gravity and *β* the gravitropic sensitivity or gain reflecting the variance in the magnitude of gravitropic responses across different species (38). The second term represents *prioprioception* which can be thought of as a counter-curving response (38), where *γ* is its gain, *R* is the radius of the organ, 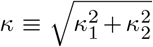is the norm of the bending curvature, and 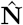is the normal director of the Frenet-Serret frame. The matrix **R**(*s, t*) in the gravitropic term represents the sensing mechanism: in the case of local sensing used here **R**(*s, t*) = **I**, and in the case of apical sensing where sensing occurs at the tip alone we set **R**(*s, t*) = **Q**(*s, t*)**Q**^*T*^ (*l*(*t*), *t*). This rotation matrix takes vectors from the material frame at the apex, *s* = *l*(*t*), to points along the growth zone *s*, and assures that the directional signal sensed at the apex is instantaneously transferred to every arc-length *s*. Results of simulations with apical sensing are described in SI Appendix 5 and SI Appendix 7. We model circumnutations as resulting from differential growth rotating around the center-line without introducing twist (30, 31, 33, 39)

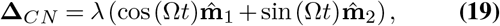

where *λ* is the circumnutation gain, Ω is its temporal angular frequency and 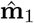 _1_ and 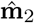 _2_ are unit vectors of a normal development of the centerline (30). We assume that the two processes are additive, allowing to take **Δ** = **Δ**_*CN*_ + **Δ**_*g*_.

### Adding a twisting material frame

Various *Arabidopsis* mutants exhibiting skewing angles also seem to exhibit a twisting cell profile (46–48), which can be interpreted as a twisting material frame around the centerline of the organ, described by the relative angle *ξ* ≡ arccos(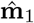 • **d**_1_) (Fig. c). Based on experimental observations, we chose a twist profile where the angle between the cell files and the root axis increases from the apex, where it is zero, to the mature zone where it reaches a constant value (Figs. b-c). This is implemented using a linear profile for the arc-length derivative of *ξ*

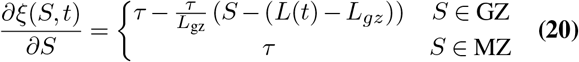

with GZ is the growth zone *L*(*t*) − *L*_*gz*_ ≤ *S* ≤ *L*(*t*), and MZ is the mature zone 0 ≤ *S < L*(*t*) − *L*_*gz*_, where *L*(*t*) is the length of the organ and *L*_gz_ is the length of the growth zone. Here *τ* represents twist, where a positive or negative value represents a right or left-handed twist respectively. The intrinsic angle between the cell files and the root axis is arctan 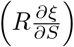 with a maximal value in the mature zone of arctan(*Rτ*). The angular frequency of rotation of the tissue at the tip follows 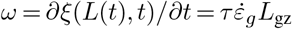 clockwise (looking from the base to the apex, see Fig. c). When incorporating twist into the dynamics of Eq. 4, only the component in the direction of **d**_3_ is affected, reading

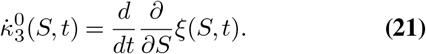

See more details in SI Appendix 2.

### Mechanics

Mechanical equilibrium is achieved at each time step when the time independent Cosserat rod equations are fulfilled along the centerline (37)

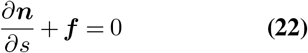

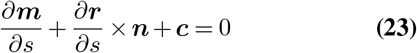

where ***n*** are the internal contact forces, ***τ*** are the internal torques or bending moments, ***f*** are the external forces per unit length and ***c*** are the external couples per unit length, all functions of *s* and *t*. The full elastic relaxation dynamics of the second step are based on the Cosserat model with a dissipation mechanism, as described in (37) and SI Appendix 3B. We follow (37) and choose a constitutive model that assumes linear elasticity, namely the internal contact forces are linearly related to the shear and stretch of the centerline, and the internal torques are linearly related to the bending and twisting

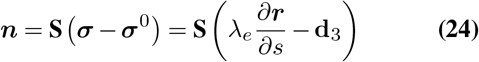

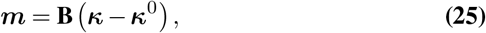

where **S** and **B** are stiffness matrices which are related to the cross sectional area and the second moment of inertia and are therefore diagonal in the local material coordinates. We also assume volume conservation, varying the local radius of the organ in order to compensate for the elastic stretch 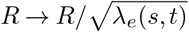.

### Simulating *Arabidopsis thaliana* roots

We simulate classic waving experiments on *Arabidopsis thaliana*, placing a rod on a plane tilted at an angle *α* with respect to the vertical (see Fig. d), allowing it to grow during 1500-3000 growth time-steps, which increases their length by a factor of 15-30. We adopt values for *Arabidopsis thaliana*, taking the radius *R* = 0.1mm (40, 58), and an initial length *L*_0_ = 1.1mm (see Tabel 1). We assume that growth is restricted to a sub-apical *growth zone* of length *L*_gz_, such that 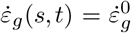 within the growth zone *L*(*t*) − *L*_*gz*_ ≤ *S* ≤ *L*(*t*), and 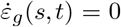 elsewhere (8) (see Fig. b and e). According to Eq. 16, the maximal growth velocity at the tip is 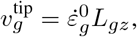, so that 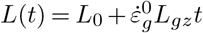. In our simulations we take *L*_*gz*_ = 1mm and 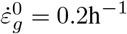 following experimental data (40). We assume the roots are incompressible with an effective Poisson’s ratio of 0.5 and Young’s modulus of *E* = 50MPa (8, 67), yielding the stiffness matrices **S** = *EπR*^2^ • diag(8*/*9, 8*/*9, 1) and **B** = *EπR*^4^ • diag(1*/*4, 1*/*4, 1*/*6), expressed in the local material frame (37). Values are summarized in Table 1. We assume the plane is frictionless. In order to emulate the stiff agar surface we assume the plane applies a local normal restoring force with a spring constant of 100kg/s^2^ and dissipative damping of the normal velocity with a dissipation coefficient of 0.01kg/s (for more details see (37)). The mature zone adheres to the substrate due to development of root hairs and lignification (8), which we emulate by using a restoring force with a spring constant of 1000kg/s^2^ which fixes in place material that exists the growth zone. Thus only the growth zone is free to change its form, while the mature zone is only free to twist around its axis. We vary the gravitropic sensitivity between 0.05 ≤ *β* ≤ 1.0 following experimental observations (40), and vary proprioceptive sensitivity in the range 0 ≤ *γ* ≤ 10 (27), though its role in roots hasn’t been established as clearly as in shoots (58).

### Solver validation

We validate our solver in SI Appendix 3 by comparing our simulations to two analytically solvable cases: (i) a clamped organ growing in the direction of an obstacle until it buckles, and (ii) the tropic response to a constant stimulus in the case of apical sensing. In SI Appendix 3 we also provide additional information regarding the numerical discretization of growth, and a numerical criterion for mechanical equilibrium. For more details about *Elastica* and the implementation of interactions between the rod and external obstacles see (37, 49).

### Estimation of intrinsic twist profile

The intrinsic twist can be evaluated by counting the number epidermal cells that cross a line tangent to the root axis, yielding the parameter CFR (cell file rotation) such that [CFR] = Number*/*length. The average projected length of one epidermal cell on the root axis is therefore 1*/*CFR. Assuming that a cell’s width is *w* ≈ 0.01mm (48), we can estimate the angle of the cell file with respect to the root axis by

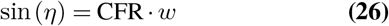

This angle in the mature zone can be expressed using *η* = arctan (*τ* · *R*), and we estimate *τ* from the CFR following

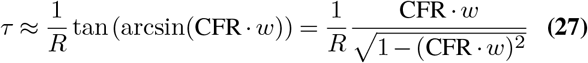

## Acknowledgements

We thank Ido Regev for helpful conversations, and for providing the experimental data from Zhang et al. (43), used in Fig.. Y.M. acknowledges funding from: Israel Science Foundation Grant (1981/14); European Union’s Horizon 2020 research and innovation program under Grant Agreement No. 824074 (Grow-Bot); Human Frontier Science Program, Reference No. RGY0078/2019. M.G. acknowledges funding from: NSF CAREER #1846752, ONR MURI N00014-19-1-2373, NSF EFRI C3 SoRo #1830881, NSF CSSI #2209322, NSF Expedition ‘Mind in Vitro’ #IIS–2123781, with computational support provided through allocation TG-MCB190004 from the Extreme Science and Engineering Discovery Environment (XSEDE; NSF grant ACI-1548562).

## Author Contributions

A.P., A.T. and Y.B. performed the research, A.P., A.T., Y.B., M.G. and Y.M. planned the research, A.P., A.T., Y.B., M.G. and Y.M. wrote the paper.

## Figure Legends

***Fig. 1*. (a) Growth patterns of *Arabidposis thaliana* roots grown on inclined agar plates**. *Arabidposis thaliana* roots are grown on inclined agar plates. While roots grow straight when the plane is vertical, at low inclinations roots tend to grow in waving patterns (17, 19, 23), and at some critical angle they coil (16, 19, 20). A number of mutants also skew at an angle ⟨*θ*⟩ (19, 60, 68). On the left, the growth patterns are shown on the scale of the whole plant (19). Green squares highlight the morphologies of interest, magnified in the middle panels, and schematically illustrated on the right. **Model definitions. (b) Cosserat rod**. Shape is defined in space with ***r***(*s, t*) along the center-line *s* at time *t*, and a local orthonormal material frame {**d**_1_(*s, t*), **d**_2_(*s, t*), **d**_3_(*s, t*)} (orange and pink lines mark ±**d**_1_(*s, t*)). Densities of external forces ***f*** (*s, t*) and couples ***c***(*s, t*) cause the rod to deform elastically. The curvature vector ***κ***(*s, t*) describes how the material frame bends and twists along the center-line, and the local vector ***σ***(*s, t*) describes how the rod shears and stretches. The light blue region near the tip represents the growth zone. **(c) Cross-section plane. d**_1_ and **d**_2_ are the material frame directors, and **d**_3_ points to the reader. 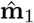 and 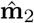 represent an arbitrary normal development of the center-line, or the *Bishop frame* (69). 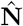 and 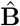 are the normal and bi-normal directors of the *Frenet-Serret* frame. The angle *ϕ* between 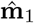 and 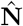 is related to the torsion of the center-line (see SI Appendix 2). Twist is described by the arc-length variations in the angle *ξ* = *ϕ* + *φ*, between 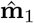 and **d**_1_. **(d) Integration scheme**. (left) Initial geometry of our simulations, where a root of length *L*_0_ is placed on a surface inclined at an angle *α* from the vertical, parameterized by an initial arc-length *S*_0_ ∈ [0, *L*_0_]. The differential growth vector related to gravitropism is parallel to the direction of gravity (see main text), and can be decomposed into normal and parallel components with respect to the plane (see SI Appendix 6). The center and right panels describe the two-step numerical integration scheme detailed in the Methods. (center) Growth-driven changes implemented to the reference configuration, following Eq. 3,4, lead to an intermediate virtual configuration disregarding elastic responses. The arc-length is parameterized by *S* ∈ [0, *L*(*t*)], related to the initial configuration through the growth stretch *λ*_*g*_(*S*_0_, *t*) = ∂*S/*∂*S*_0_. (right) Relaxation to mechanical equilibrium following Eqs. 1 and 2. The actual configuration of the rod is parameterized by *s* ∈ [0, *l*(*t*)], related to the virtual configuration through the elastic stretch *λ*_*e*_(*S, t*) = ∂*s/*∂*S*. **(e) Roots axial growth profile**. Micrograph of an *Arabidopsis thaliana* root. Cells divide at the tip, and elongate within the *growth zone*, until they stop elongating, reaching the *mature zone*. Image courtesy of Eilon Shani.

***Fig. 2*. Simulations reproduce waving and coiling. (a) Final configurations**. Top view of final configurations of simulations for tilt angles 0^0^ ≤ *α* ≤ 90^0^, with *β* = *γ* = 0.1. Pink and orange lines represent the direction of the vectors ±**d**_1_ describing the material frame along the organ. The local planar angle *θ*(*s, t*) is the angle between the projection of the organ tangent on the tilted plane, and the projection of the direction of gravity **ĝ**_∥_. **(b) Snapshots of waving dynamics**. Five snapshots during the developing waving pattern for a root at *α* = 40^*o*^, with local gravisensing, *β* = 0.2, and *γ* = 0.1. Stages (i)-(ii) depict the first symmetry breaking due to an elastic instability that reduces bending energy by twisting sideways. Between stages (iii) and (v) left-handed twist is being accumulated in the mature zone as the root turns clock-wise (evident by following **d**_1_), in line with observed twisting cell files (26). **(c)**-**(e): Properties of the growth zone during the development of a waving pattern**, corresponding to the root in (b). Dashed lines represent the snapshots in (b), and time is shown in number of time-steps. **(c)** Planar tip angle *θ*^tip^(*t*) (blue). For comparison, we show the behavior of a corresponding free organ (pink line), that is of a root growing without substrate but characterized by the same initial angle (with respect to gravity) of the waving root (blue line). Both profiles present oscillations with a similar period, however in the absence of interactions with the plane the oscillations decay in time. **(d)** Average elastic bending energy *E*_*b*_ and twisting energy *E*_*t*_ in the growth zone. **(e)** Total writhe Wr and twist Tw in the growth zone.

***Fig. 3*. Comparison of model predictions with simulations and experiments. (a) Curvature of waving patterns originate from the gravitropic response of a free organ**. The maximal curvature max *κ* of the projected shape of simulated waving patterns on the plane approximately scales like the predicted maximal curvature of a free organ 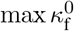 (Eq. 8), tilted at the same angle *α*, as suggested by Eq. 6. Simulations were run with *L*_gz_*/R* = 10, *γ* = 0.1, and 0.05 ≤ *β* ≤ 1.0 (see Table 1), and for a range of inclinations *α*. Results for larger values of *γ* and apical gravisensing are brought in SI Appendix 5. Dashed line represents identity, for reference. **(b) Wavelength vs. amplitude**. Following our proposed scaling laws for the wavelength *λ* and amplitude *A* of waving patterns (Eqs. 9,10,11), *λ* is plotted against 4*A* cot(*α*) for values measured in simulations and experiments (16, 43, 59), exhibiting a linear dependence as predicted. Dashed line represents identity, for a reference. The wavelength and amplitude of each simulation are quantified by averaging the distances between neighboring peaks of the waving pattern parallel and normal to the downhill direction. Data points for *α* ≤ 10 deg from (43) are not shown as they diverge. **(c) Geometric relation between wavelength and amplitude**. The angle on the plane is the integral over curvature, and following (a) is dictated by the tip angle of a free organ, which can be approximated by the inclination angle *α* (see virtual configuration in Fig. e). **(d) Configuration space**. Shapes of simulated organs (with *L*_*gz*_*/R* = 10 and *γ* = 0.1) for different values of the effective gravitropic gain *βL*_gz_*/R* and tilt angle *α*. Symbols represent the final configuration shape of simulations (◼ symbols for straight, ♦ for waving,• for coiling). Simulations agree with model predictions, represented by background color: transitions from straight to waving occur at critical angle *α*_s→w_ (Eq. 12), and transitions to coiling occur at *α*_w→c_ (Eq. 13), estimated in SI Appendix 7. The second column represents the configurations shown in Fig. a. Equivalent configuration spaces for high *γ* and apical sensing in SI Appendix 7. **(e) Observed coiling transition**. Experimental observations of coiling probability (43) agree with model prediction (represented by background colors as in (e) for *βL*_gz_*/R* = 0.5). We note that (43) reports probabilities only for the emergence of coiling behavior, with the white portion of the plot encompassing the combined probability of observing straight or waving patterns.

***Fig. 4*. Simulations and model reproduce skewing patterns. (a) Simulation configurations**. Top view of simulated organs for different twisting frequencies *ω* and a constant turning time *T*_0_ (with *α* = 30^*o*^, *γ* = 0.1, and *β* = 0.2). **(b) Observed cell file rotation** of an Arabidopsis mutant exhibiting skewing, manifesting intrinsic twist (47). **(c) Model for intrinsic twist**. The angle between the cell files and the root axis increases from the apex to the mature zone, captured by the angular velocity *ω*, as described by Eq. 5. **(d)**-**(f) Properties of the growth zone during the development of a skewed pattern**. Similar to Fig. (c)-(e), for *ωT*_0_ = 1. Symmetry is broken by twist, without an elastic instability. A comparison between various values of *ωT*_0_ appears in SI Appendix, Fig. S10. **(g) Relation between skewing angles** ⟨*θ*⟩ **and intrinsic twist**. Following the limiting relations in Eqs. 14 and 15 we plot sin ⟨*θ*⟩*/* tan *α* vs. *ωT*_0_ • *R/L*_*gz*_, for both simulations and experiments (26, 60–62). Simulations were run with *L*_gz_*/R* = 10, *γ* = 0.1, 0.1 ≤ *β* ≤ 0.25, 10^*o*^ ≤ *α* ≤ 40^*o*^, and 0 ≤ *Rτ* ≤ 0.8, which give 0 ≤ *ωT*_0_ ≤ 9.3 (see Table 1). Experimental data (26, 60–62) are plotted assuming *L*_gz_*β/R* = 0.5, and *ω* is estimated from experimental values of cell file rotation (CFR) via Eqs. 27 and Eq. 5. Predicted asymptotic relations in the limits *ωT*_0_ ≪ 1 and *ωT*_0_ ≫ 1, as detailed in Eqs. 14 and 15, are represented by solid lines. Experiments agree with simulations and asymptotic predictions, except for MBD-GFP mutants (26). In SI Appendix, Fig. S9, we plot the correlation between intrinsic twist *τ* and skewing angle ⟨*θ*⟩ for the experimental data.

