## Supplementary material for "On the mechanical origins of waving, coiling and skewing in *Arabidopsis thaliana* roots"

### – Supplementary Information –
### On the mechanical origins of waving, coiling and skewing growth patterns in plant roots

Amir Porat[a],Arman Tekinalp[b],Yashraj Bhosale[b],Mattia Gazzola[b],Yasmine Meroz[c]

[a]Faculty of Exact Sciences, School of Physics, Tel Aviv University, Tel Aviv, Israel
[b]Mechanical Sciences and Engineering, University of Illinois at Urbana–Champaign, Urbana, IL 61801, USA
[c]Faculty of Life Sciences, School of Plant Sciences and Food Security, Tel Aviv University, Tel Aviv, Israel

**8 Circumnutations** **14**

**9 Skewing - additional figures** **14**

### 1. Captions for supplementary movies

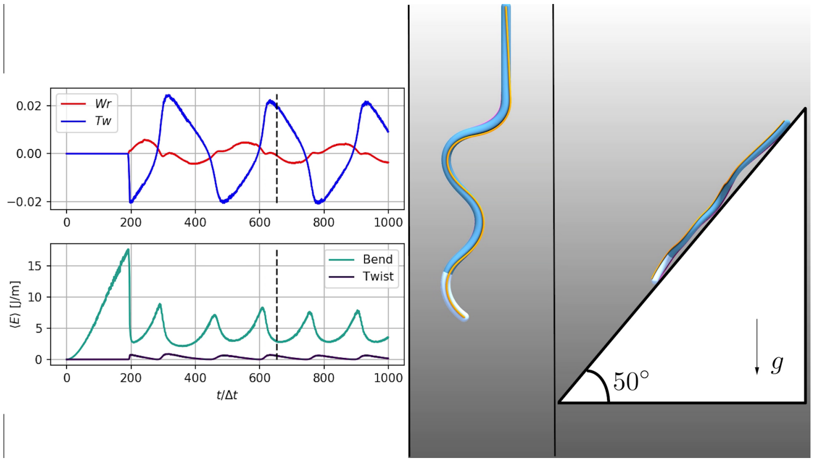

**Movie S1.** Development of a waving pattern for a simulated root with $\alpha = 40^o$, local gravisensing, $\beta = 0.2$, and $\gamma = 0.1$. The movie presents both a side view and a top view normal to the tilted plane. Next to the movie we present the development of geometric and elastic parameters of the growth zone, as described in Fig. 2b,d,e.

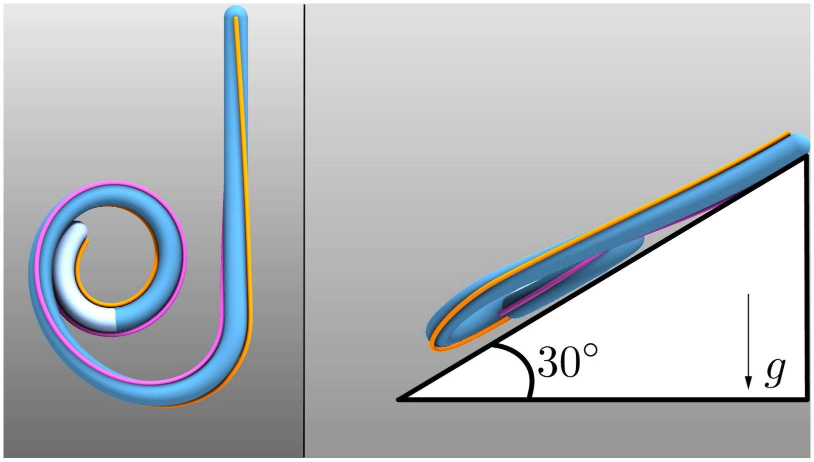

**Movie S2.** Development of a coiling pattern for a simulated root with $\alpha = 60^o$, local gravisensing, $\beta = 0.1$, and $\gamma = 0.1$ (as in Fig. 2a). The movie presents both a side view and a top view normal to the tilted plane.

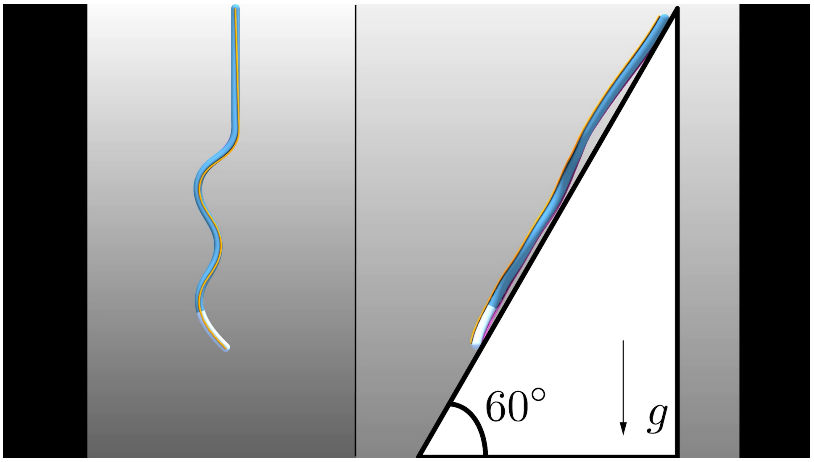

**Movie S3.** Development of a waving patterns for simulated roots with $\alpha = 30^o$, local gravisensing, $\beta = 0.2$, and $\gamma = 0.1$. Here $\omega T_0 = 0$, while supplementary movies 4 and 5 present skewing with $\omega T_0 = 1$ and $\omega T_0 = 8.3$ correspondingly (as in Fig. 4a and Fig. S10). The movie presents both a side view and a top view normal to the tilted plane.

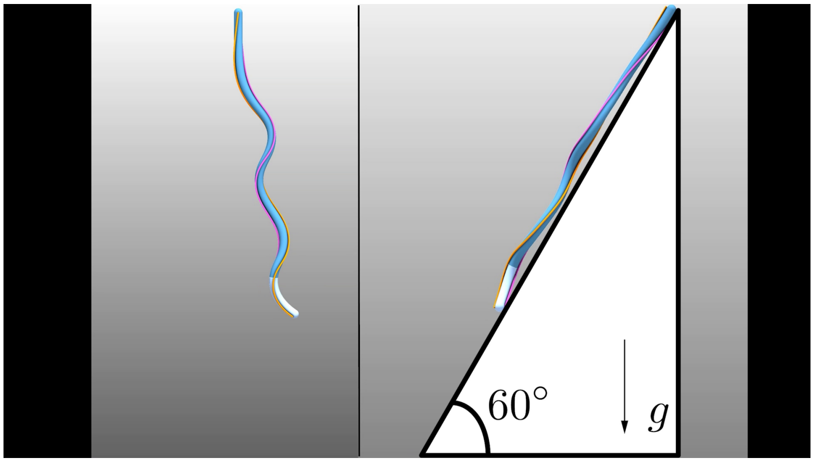

**Movie S4.** Development of a skewing patterns for simulated roots with $\alpha = 30^o$, local gravisensing, $\beta = 0.2$, and $\gamma = 0.1$. Here $\omega T_0 = 1$, while supplementary movies 3 and 5 present skewing with $\omega T_0 = 0$ and $\omega T_0 = 8.3$ correspondingly (as in Fig. 4a and Fig. S10). The movie presents both a side view and a top view normal to the tilted plane.

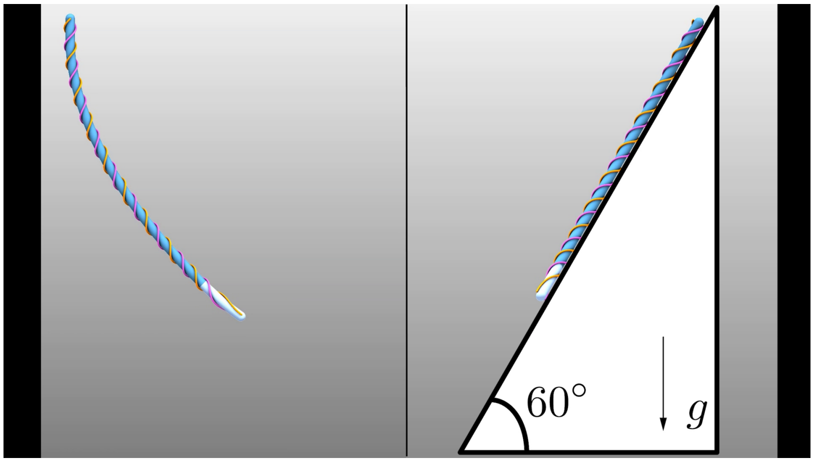

**Movie S5.** Development of a skewing patterns for simulated roots with $\alpha = 30^o$, local gravisensing, $\beta = 0.2$, and $\gamma = 0.1$. Here $\omega T_0 = 8.3$, while supplementary movies 3 and 4 present skewing with $\omega T_0 = 0$ and $\omega T_0 = 1$ correspondingly (as in Fig. 4a and Fig. S10). The movie presents both a side view and a top view normal to the tilted plane.

#### 2. Growth model

We here develop Eq. 4 from the main article. We assume for simplicity that no external forces act on the rod such that $s = S$ and $\boldsymbol{\kappa} = \boldsymbol{\kappa}^0$. The curvature vector of a center-line that has an intrinsic twist profile can be described in the local material frame as (2):

$$\boldsymbol{\kappa}^0 = \kappa_1^0 \mathbf{d}_1 + \kappa_2^0 \mathbf{d}_2 + \kappa_3^0 \mathbf{d}_3 = \kappa \sin(\varphi) \mathbf{d}_1 + \kappa \cos(\varphi) \mathbf{d}_2 + \left(\tau + \frac{\partial \varphi}{\partial s}\right) \mathbf{d}_3, \qquad \text{[S1]}$$

where $\kappa = \sqrt{(\kappa_1^0)^2 + (\kappa_2^0)^2}$ is the bending curvature, $\tau$ is the torsion and $\varphi$ is the register angle between the normal director of the local Frenet-Serret frame $\hat{\mathbf{N}}$ and the material frame's director $\mathbf{d}_1$ (see Fig. 1c), all functions of arc-length and time. We note that the torsion can be described using a register angle $\phi$ between $\hat{\mathbf{N}}$ and a director from the normal development of the center-line $\hat{\mathbf{m}}_1$, such that $\frac{\partial \phi}{\partial s} = \tau$.

To describe an intrinsic twist profile independently from the normal director $\hat{\mathbf{N}}$, we use the register angle between $\hat{\mathbf{m}}_1$ and $\mathbf{d}_1$, and mark it as $\xi = \phi + \varphi$. We assume that an arbitrary twist profile doesn't vary the shape of the center-line in space, only the orientation of the material frame along it via $\varphi$.

The time derivative of the components of the intrinsic curvature vector from Eq.S1 gives:

$$\dot{\kappa}_1^0 = \dot{\kappa} \sin(\varphi) + \kappa \cos(\varphi) \dot{\varphi} \qquad \text{[S2]}$$

$$\dot{\kappa}_2^0 = \dot{\kappa} \cos(\varphi) - \kappa \sin(\varphi) \dot{\varphi} \qquad \text{[S3]}$$

$$\dot{\kappa}_3^0 = \dot{\tau} + \frac{\partial \dot{\varphi}}{\partial s} = \frac{\partial \dot{\phi}}{\partial s} + \frac{\partial \dot{\varphi}}{\partial s} = \frac{\partial \dot{\xi}}{\partial s} \qquad \text{[S4]}$$

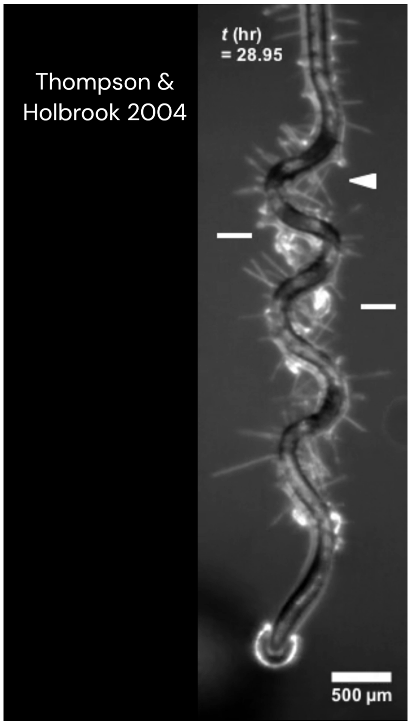

**Movie S6. Time-lapse of a developing waving pattern from (1), reproduced here for reference.**

where $\dot{x} = \frac{\partial x}{\partial t} + v_g \frac{\partial x}{\partial s} \equiv \frac{Dx}{Dt}$ is a material derivative which accounts for the growth of the centerline, and is calculated with respect to the rotating material frame.
To continue, we quote the growth driven dynamics of the centerline's intrinsic curvature as was derived in (3) and in (4) for a cylindrical organ with a circular cross-section of radius $R$:

$$\dot{\kappa} = \frac{\dot{\varepsilon}_g}{R} \mathbf{\Delta} \cdot \hat{\mathbf{N}} \quad \text{[S5]}$$

$$\dot{\phi} = \frac{\dot{\varepsilon}_g}{R\kappa} \mathbf{\Delta} \cdot \hat{\mathbf{B}} \quad \text{[S6]}$$

here $\mathbf{\Delta}$ is the differential growth vector, which in the case of gravitropic roots defined by Eq. 3 in the main text. Plugging Eqs. S5-S6 in Eqs. S2-S3 and using $\dot{\varphi} = \dot{\xi} - \dot{\phi}$ gives:

$$\dot{\kappa}_1^0 = \frac{\dot{\varepsilon}_g}{R} \mathbf{\Delta} \cdot \left( \hat{\mathbf{N}} \sin(\varphi) - \hat{\mathbf{B}} \cos(\varphi) \right) + \kappa \cos(\varphi) \dot{\xi} \quad \text{[S7]}$$

$$\dot{\kappa}_2^0 = \frac{\dot{\varepsilon}_g}{R} \mathbf{\Delta} \cdot \left( \hat{\mathbf{N}} \cos(\varphi) + \hat{\mathbf{B}} \sin(\varphi) \right) - \kappa \sin(\varphi) \dot{\xi} \quad \text{[S8]}$$

Noticing that $\mathbf{d}_2 = -\hat{\mathbf{N}} \sin(\varphi) + \hat{\mathbf{B}} \cos(\varphi) = \mathbf{d}_3 \times \mathbf{d}_1$ and that $\mathbf{d}_1 = \hat{\mathbf{N}} \cos(\varphi) + \hat{\mathbf{B}} \sin(\varphi) = \mathbf{d}_2 \times \mathbf{d}_3$ then gives:

$$\dot{\kappa}_1^0 = \frac{\dot{\varepsilon}_g}{R} (\mathbf{d}_3 \times \mathbf{\Delta}) \cdot \mathbf{d}_1 + \kappa_2^0 \dot{\xi} \quad \text{[S9]}$$

$$\dot{\kappa}_2^0 = \frac{\dot{\varepsilon}_g}{R} (\mathbf{d}_3 \times \mathbf{\Delta}) \cdot \mathbf{d}_2 - \kappa_1^0 \dot{\xi} \quad \text{[S10]}$$

$$\dot{\kappa}_3^0 = \frac{\partial \dot{\xi}}{\partial s} \quad \text{[S11]}$$

These dynamics can be written in the vector form using a material time derivative, written in the material frame:

$$\dot{\boldsymbol{\kappa}}^0 = \dot{\kappa}_1^0 \mathbf{d}_1 + \dot{\kappa}_2^0 \mathbf{d}_2 + \dot{\kappa}_3^0 \mathbf{d}_3 = \frac{\dot{\varepsilon}_g}{R} (\mathbf{d}_3 \times \mathbf{\Delta}) - \dot{\xi} \mathbf{d}_3 \times \boldsymbol{\kappa}^0 + \frac{\partial \dot{\xi}}{\partial s} \mathbf{d}_3 \quad \text{[S12]}$$

The dynamics of an organ with no intrinsic twist ($\xi = 0$, $\kappa_3^0 = 0$) can thus be written using:

$$\dot{\boldsymbol{\kappa}}^0 = \dot{\kappa}_1^0 \mathbf{d}_1 + \dot{\kappa}_2^0 \mathbf{d}_2 = \frac{\dot{\varepsilon}_g}{R} (\mathbf{d}_3 \times \mathbf{\Delta}) \quad \text{[S13]}$$

which gives the Eq. 4 in the main article. Moreover, following Eq. S12, a constant center-line in space ($\dot{\varepsilon}_g = 0$) with a time varying register angle $\xi$ which is unifrom in arc-length $\frac{\partial \xi}{\partial s} = 0$ changes the curvature vector according to:

$$\dot{\boldsymbol{\kappa}}^0 = -\dot{\xi}\mathbf{d}_3 \times \boldsymbol{\kappa}^0 \quad \text{[S14]}$$

Eq. S14 shows that the bending curvature performs a contravariant rotation which keeps it directed to a constant direction in space as the register angle $\xi$ varies in time. To rotate the bending curvature with the rotation of the material frame as occurs in plants, we now assume that temporal variations in $\xi$ do affect the shape of the centerline by omitting the contravariant rotation from Eq. S12:

$$\dot{\boldsymbol{\kappa}}^0 = \frac{\dot{\varepsilon}_g}{R}\left(\mathbf{d}_3 \times \boldsymbol{\Delta}\right) + \frac{\partial \dot{\xi}}{\partial s}\mathbf{d}_3 \quad \text{[S15]}$$

which gives Eq. 21 from the methods section. This relation can also be derived by changing Eq. S6 to:

$$\dot{\phi} = \frac{\dot{\varepsilon}_g}{R\kappa}\boldsymbol{\Delta} \cdot \hat{\mathbf{B}} + \dot{\xi} \quad \text{[S16]}$$

and repeating the derivation. Equation S16 treats the variations of intrinsic twist $\dot{\xi}$ as a source of torsion which is added to the torsion caused by differential growth, and substituting it in $\dot{\varphi} = \dot{\xi} - \dot{\phi}$ shows that $\varphi$ varies only due to differential growth.

#### 3. Discretization and validation of the numerical solver

**A. Growth discretization.** Our solver is based on Elastica, which is presented in (5). The solver discretizes a Cosserat rod using a chain of $n+1$ point masses located on vertices. The arc-length of the rod is then discretized to $n$ straight edges connecting the masses with lengths $\ell_i$, where $i \in \{1, .., n\}$. The full state of the rod is described using variables located on the vertices ($i \in \{1, .., n+1\}$), variables of the edges between them ($i \in \{1, .., n\}$), and variable located only on the interior vertices ($i \in \{1, .., n-1\}$), which are the average of the integrated values over Voronoi domains. The length of a Voronoi domain surrounding mass $i+1$ is defined by:

$$\mathcal{D}_i = \frac{\ell_i + \ell_{i+1}}{2} \quad \text{[S17]}$$

Here, we illustrate how we input the growth dynamics described in (4) into the discretized rod. We use variable symbols directly from (5), and keep the convention of putting a hat over variables in the reference stress-free configuration.
Given a growth law by the smooth functions $\dot{\varepsilon}_g$ and $\boldsymbol{\Delta}$, we begin by discretizing the relative growth rate for each reference edge $\hat{\ell}_i$ using an average over its length:

$$\dot{\varepsilon}_{g,i}(t) = \frac{1}{\hat{\ell}_i}\int_{S_i}^{S_{i+1}} \dot{\varepsilon}_g(S, t)dS \quad \text{[S18]}$$

where $S$ is the arc-length of the reference configuration and $\hat{\ell}_i = S_{i+1} - S_i$ is the length of edge $i$. By definition, the incremental elongation of edge $i$ is related to the relative growth rate according to:

$$\dot{\varepsilon}_g = \frac{1}{\hat{\ell}}\frac{d\hat{\ell}}{dt} \quad \text{[S19]}$$

Assuming a growth time-step $\Delta t$ then gives a discretized elongation accroding to:

$$\hat{\ell}_i(t + \Delta t) = (1 + \dot{\varepsilon}_{g,i}(t)\Delta t)\hat{\ell}_i(t), \quad \text{[S20]}$$

where $\dot{\varepsilon}_{g,i}(t)\Delta t$ is an incremental growth strain. Similar to the rest length $\hat{\ell}_i$, many other properties of the rod must be updated, such as an edge's volume $\hat{V}_i$ and the mass second moment of inertia $\hat{J}_i$, both proportional to the edge's reference length $\hat{\ell}_i$. Growth processes also increase the mass of the grown edge, which is divided between the adjacent vertices. This mass increase can be performed using the operator $\mathcal{A}^h$ presented in (5):

$$m_i(t + \Delta t) = m_i(t) + \mathcal{A}^h\left(m_i(t)\dot{\varepsilon}_{g,i}\right)\Delta t. \quad \text{[S21]}$$

Having discretized the relative growth rate, we turn to discretize the dynamics of intrinsic curvature from Eqs. S9-S10. Since each discretized edge flows with growth, the discretization index $i$ refers to the Lagrangian coordinate, and we can use Eqs. S9-S10 directly. In (5), the discretization of the smooth curvature vector is done by assigning each interior vertex with the average of the curvature vector over its Voronoi domain:

$$\boldsymbol{\kappa}_i^0(t) = \frac{1}{\hat{\mathcal{D}}_i}\int_{S_i - \hat{\ell}_i/2}^{S_i + \hat{\ell}_{i+1}/2} \boldsymbol{\kappa}^0(S, t)dS \equiv \left\langle \boldsymbol{\kappa}^0(t)\right\rangle_i \quad \text{[S22]}$$

where we marked the $i$-th average using $< \cdot >_i$. Taking a time derivative of Eq. S22 gives:

$$\dot{\boldsymbol{\kappa}}_i^0(t) = -\frac{\dot{\hat{\mathcal{D}}}_i}{\hat{\mathcal{D}}_i}\boldsymbol{\kappa}_i^0(t) + \frac{1}{2\hat{\mathcal{D}}_i}\left(\dot{\hat{\ell}}_i\hat{\boldsymbol{\kappa}}^0(S_i - \hat{\ell}_i/2, t) + \dot{\hat{\ell}}_{i+1}\hat{\boldsymbol{\kappa}}^0(S_i + \hat{\ell}_{i+1}/2, t)\right) + \left\langle \dot{\boldsymbol{\kappa}}^0(t)\right\rangle_i \quad \text{[S23]}$$

where the first term is the time derivative of the Voronoi domain, the second term are the derivatives of the limits of the integral, and the last term is the derivative of the integrand. We assume that the discretization is fine, such that the curvature changes slowly from one edge to another (i.e., $\boldsymbol{\kappa}_i^0(t) \approx \hat{\boldsymbol{\kappa}}^0(S_i \pm \hat{\ell}_i/2, t)$). Then, the first two terms cancel each other, and the main contribution comes from the last term. In the material frame, inserting Eq. S13 into Eq. S23 while omitting the time dependence for brevity gives (where $\mu \in \{1, 2\}$):

$$\dot{\kappa}_{\mu,i}^0 \approx \left\langle \dot{\kappa}_\mu^0 \right\rangle_i = \left\langle \frac{\dot{\varepsilon}_g}{R} (\mathbf{d}_3 \times \mathbf{\Delta}) \cdot \mathbf{d}_\mu \right\rangle_i \tag{S24}$$

The average in the last term in Eq. S24 depends on several non-trivial continuous fields, and is difficult to solve for any arbitrary growth rate or differential growth vector. We therefore approximate it by using it's value on the interior vertex, which we mark using the notation *int*:

$$\dot{\kappa}_{\mu,i}^{0,\text{int}} = \frac{1}{R} \frac{\dot{\varepsilon}_{g,i}\hat{\ell}_i + \dot{\varepsilon}_{g,i+1}\hat{\ell}_{i+1}}{\hat{\ell}_i + \hat{\ell}_{i+1}} (\mathbf{d}_{3,i}^{\text{int}} \times \mathbf{\Delta}_i^{\text{int}}) \cdot \mathbf{d}_{\mu,i}^{\text{int}}. \tag{S25}$$

In (5), the discretized coordinate frame $\boldsymbol{Q}_i = \{\mathbf{d}_{1,i}, \mathbf{d}_{2,i}, \mathbf{d}_{3,i}\}^{-1}$ describes the frame at the center of edge $i$. To find the frame on the interior vertex which is used in Eq. S25, $\boldsymbol{Q}_i$ is propagated to the location of vertex $i+1$ using a rotation matrix expressed by the exponential map (4):

$$\boldsymbol{Q}_i^{\text{int}} = \boldsymbol{Q}_i \exp\left( -\frac{\hat{\ell}_i}{2} \begin{pmatrix} 0 & -\kappa_{i,3}^0 & \kappa_{i,2}^0 \\ \kappa_{i,3}^0 & 0 & -\kappa_{i,1}^0 \\ -\kappa_{i,2}^0 & \kappa_{i,1}^0 & 0 \end{pmatrix} \right) \tag{S26}$$

Lastly, iterating many growth steps may lead to very large edge lengths. To allow for long simulations while keeping the validity of the discretization, we implemented a procedure that divides an edge to two distinct edges of equal lengths when its length has grown above a given threshold. In our simulations we discretize the arc-length initially to $\hat{\ell}_i = R/4$ and divide the edges when they surpassed $R/3$. In addition, to reduce numerical errors and obtain a smooth quasi-static integration, we make sure that the total growth induced extension in one growth time-step is smaller than the numerical discretization length of the arc-length of the root. In our simulations, where we use a uniform growth rate $\dot{\varepsilon}_g^0$ and a growth zone of length $L_{gz}$, this is achieved by setting the incremetnal growth strain to be $\Delta\dot{\varepsilon}_g^0 t = 0.01$ (the total growth induced extension is $\delta L = \dot{\varepsilon}_g^0 \Delta t L_{gz}$ such that $\delta L/R = 0.1$ for $L_{gz} = 10R$). We therefore present time in the main article in units of $\Delta t$, such that it takes $100\Delta t$ to grow one growth zone length ($\dot{\varepsilon}_g^0 = 0.2\text{h}^{-1}$ gives $\Delta t = 3\text{min}$, much longer than typical elastic relaxation times).

**B. Mechanical relaxation.** Once the rest lengths and intrinsic curvatures of a rod have been updated according to a prescribed growth law after each growth timestep $\Delta t$, the rod is relaxed mechanically using smaller timesteps $\delta t$. The relaxation is preformed according to a full Cosserat rod dynamics as described in detail in (5), where the numerical method is presented and validated extensively by comparing with analytical solutions of benchmark problems. There, a dissipation model is introduced by adding friction-like terms to the external body force and couple:

$$\boldsymbol{f_\nu} = -\nu_d \frac{\partial \boldsymbol{r}}{\partial t}, \quad \boldsymbol{c_\nu} = -\nu_d \boldsymbol{\omega}, \tag{S27}$$

where $\boldsymbol{\omega}$ is the angular velocity of the local material frame such that for $i \in \{1, 2, 3\}$: $\partial \mathbf{d}_i / \partial t = \boldsymbol{\omega} \times \mathbf{d}_i$.
In our simulations we use $\nu_d = 0.25\text{kg/s}$ and a mass density of $10^4\text{kg/m}^3$. Even though this density is higher than that of plant roots, it has no affect on the quasi-static configuration, and it reduces the elastic oscillations of the rod during the relaxation dynamics (gravitational body forces can be scaled accordingly). Moreover, we use a relaxation time step that is related to the elastic shear wave speed within the organ: $\delta t \propto \ell/\sqrt{E/\rho} \sim 0.5\ \mu\text{s}$.
Since the dissipation dynamics described in Eq. S27 lead to an exponential decay of the energy, a criterion is required to determine if the system has reached mechanical equilibrium. In all of our simulations this criterion was chosen to be the stabilization of the elastic strain $\varepsilon_e$. Specifically, we assumed the system is in mechanical equilibrium if the axial elastic strain $\varepsilon_e$ changes less than $10^{-5}$ in 12500 relaxation time-steps $\delta t$ ($\sim 6$ ms).

**C. Validation.** To validate the accuracy of our 2 step integration scheme, we compare results from simulations to two analytic results regarding morpho-elastic rods.

***C.1. Case 1: Growth induced buckling.*** In this first example, we place a cylindrical rod of length $L_0$ and radius $R_0$ which is clamped at its base and connected to a spherical joint at its apex. The joint allows the apex to change direction freely, but applies a restoring force on its displacements with a spring constant $k_{\text{spring}}$. We let the rod grow with a uniform relative growth rate $\dot{\varepsilon}_g^0$ and measure the force it exerts on the spherical hinge. After an initial growth and compression, the force the rod applies on the spring surpasses the Euler critical load and the rod buckles. However, until that time the dynamics are axis-symmetric, which allows us to describe the quasi-static dynamics analytically in one dimension.
To begin, we describe the rest length $L$ at each time using the uniform relative growth rate:

$$\frac{dL}{dt} = \dot{\varepsilon}_g^0 L \longrightarrow L(t) = L_0 \exp\left(\dot{\varepsilon}_g^0 t\right) \tag{S28}$$

This is known as an exponential growth profile. Since the rod is uniform, the growth stretch is independent of arc-length and is given by:

$$\lambda_g(t) = \frac{L(t)}{L_0} = \exp\left(\dot{\varepsilon}_g^0 t\right). \tag{S29}$$

In addition, since the dynamics are quasi-static, there exists a force balance between the rod and the hinge at all times. The restoring force is spring-like and depends on the displacement of the apex:

$$\boldsymbol{F}_{\text{obstacle}}(t) = -k_{\text{spring}}(l(t) - L_0)\hat{\mathbf{T}} = -k_{\text{spring}}L_0(\lambda_e(t)\lambda_g(t) - 1)\hat{\mathbf{T}} \tag{S30}$$

where $\lambda_e(t) = l(t)/L(t)$ is the elastic stretch and $\hat{\mathbf{T}}$ is the tangent to the rod. Assuming the rod has the same constitutive equation as in the main text allows us to write the force exerted on the hinge by the rod:

$$\boldsymbol{F}_{\text{rod}}(t) = -\boldsymbol{n}(t) = -\pi\left(\frac{R_0}{\sqrt{\lambda_e(t)}}\right)^2 E(\lambda_e(t) - 1)\hat{\mathbf{T}} = \pi R_0^2 E\left(\frac{1}{\lambda_e(t)} - 1\right)\hat{\mathbf{T}} \tag{S31}$$

A force balance $\boldsymbol{F}_{\text{obstacle}}(t) = -\boldsymbol{F}_{\text{rod}}(t)$ gives an equation that couples the elastic and growth stretches:

$$\pi R_0^2 E\left(\frac{1}{\lambda_e(t)} - 1\right) = k_{\text{spring}}L_0(\lambda_e(t)\lambda_g(t) - 1) \tag{S32}$$

Solving Eq.S32 for $\lambda_e(t)$ gives:

$$\lambda_e(t) = \frac{1}{2\lambda_g(t)}\left(1 - \eta + \sqrt{(1-\eta)^2 + 4\eta\lambda_g(t)}\right) \tag{S33}$$

where $\eta = \pi E R_0^2/(k_{\text{spring}}L_0)$. Plugging Eqs. S29 and S33 into Eq.S31 gives a force profile as a function of time. This profile is valid as long as the force is smaller than the Euler critical load:

$$F_{cr}(t) = \frac{\pi^2 EI(t)}{(Kl(t))^2} == \frac{\pi^3 ER^4(t)}{(2Kl(t))^2} = \frac{\pi^3 ER_0^4}{(2K\lambda_e^2(t)\lambda_g(t)L_0)^2} \tag{S34}$$

where $I = \pi R^4/4$ is the second moment of area and $K$ is a dimensionless constant which depends on the boundary conditions known as the effective length factor (6).

We simulate these dynamics using a rod with the same parameters as in the main article, except all the lengths are multiplied by 10, and the following parameters are changed: $\ell_0 = R_0/6$, $\dot{\varepsilon}_g^0 = 2.5 \cdot 10^{-3}/\Delta T$, $\beta = 0$ and $\gamma = 0$. We take a hinge with a spring constant of $k_{\text{spring}} = 10^4 \text{kg}/\text{sec}^2$ and add a gravitational forced with a gravity vector that is tilted by $1^o$ degree from the initial tangent to break the symmetry. The theoretical and simulated force as a function of time are plotted in figure S1, as well as the buckling point, with an excellent agreement.

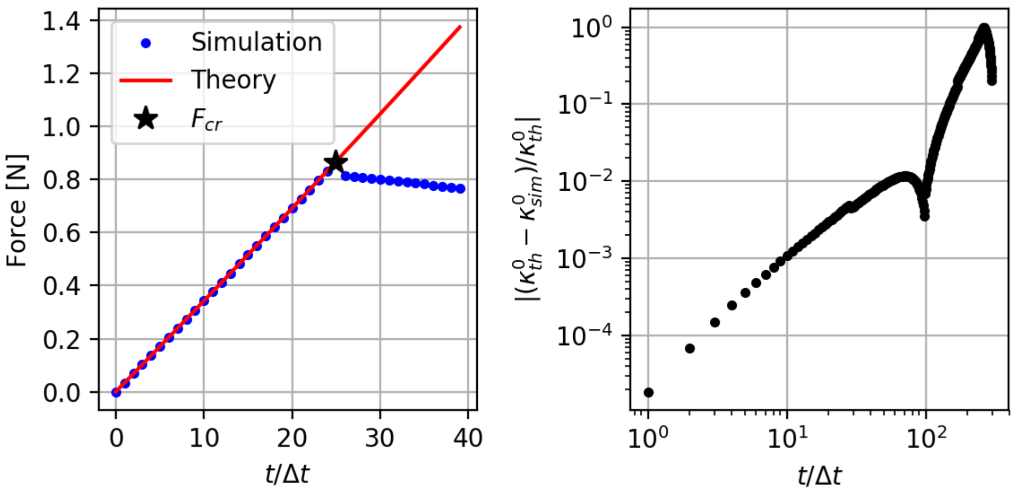

**Fig. S1. Solver validation.** Left: Growth induced buckling. Comparison between the theoretical force the rod exerts on the hinge as a function in time to the results from a corresponding simulation, including the buckling of the growing rod. Right: The error in the simulated curvature of the growth zone during a tropic movement of an organ with apical sensing and a sub-apical growth zone.

***C.2. Case 2: Tropic movement with apical sensing and a sub-apical growth zone.*** To validate the discretization of dynamics of the curvature vector, we compare our simulation to a 2-d analysis of growing organs presented in (7). There, gravitropic roots are modeled using a uniform sub-apical growth zone and apical gravisensing, in a similar fashion to the modeling used here. There, the curvature in the growth zone is uniform and obeys the following dynamics:

$$-\frac{(y''+\gamma y')}{\sqrt{\eta^2-(y'+\gamma y)^2}} = y + y' \qquad \text{[S35]}$$

where $y = L_{gz}\kappa^0$, $y' = \frac{dy}{dx}$, $x = \dot{\varepsilon}_g t$, $\eta = \beta L_{gz}/R$ and $\gamma$ is the proprioceptive gain. We compare the analytical solution of Eq. S35 to the curvature from a simulation of an organ with the the same parameters in the main article and $\beta = 0.1$, $\gamma = 0$, and an initial tilt of $90^o$ with respect to gravity. We quantify the error in the simulated intrinsic curvature by $|(\kappa^0_{\mathrm{th}} - \kappa^0_{\mathrm{exp}})/\kappa^0_{\mathrm{th}}|$, where $\kappa^0_{\mathrm{th}}$ is the solution of Eq. S35 and $\kappa^0_{\mathrm{exp}}$ is the simulated intrinsic curvature at the apex. As shown in Fig. S1, for the first $\sim 100$ growth time-steps the error is below 1 percent.

#### 4. Writhe and twist

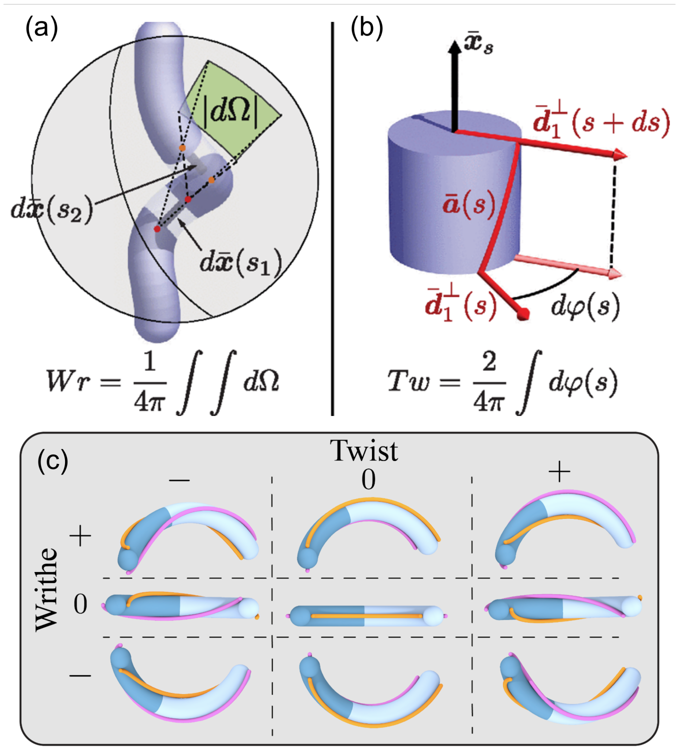

**Fig. S2. Writhe and Twist.** (a) Writhe (Wr) equals the centerline's average oriented self-crossing number computed in terms of the integral of the solid angle $d\Omega$ determined by the infinitesimal centerline segments $d\bar{\boldsymbol{x}}(s_1)$ and $d\bar{\boldsymbol{x}}(s_2)$ (left-handed intersections are negative). (b) Twist (Tw) is the integral of the infinitesimal rotations $d\varphi$ of the auxiliary curve $\bar{\boldsymbol{a}}$ around $\bar{\boldsymbol{x}}_s$. Here, the vector $\bar{\boldsymbol{a}}$ traced out by $\bar{\boldsymbol{d}}_1^\perp$ (i.e., the projection of $\bar{\boldsymbol{d}}_1$ onto the normal-binormal plane) is shown in red. For a closed curve Lk = Tw + Wr, where Lk (link) is the average oriented crossing number of $\bar{\boldsymbol{x}}(s)$ with $\bar{\boldsymbol{a}}(s)$. (c) Illustration of the difference between writhe and twist: Writhe describes 3D rotations of the center-line and is being used here as a measure of the torsion of the growth zone's center-line, being 0 when the center-line can be embedded on a plane, positive when it resembles a right-handed helix and negative when it resembles a left-handed helix. In contrast, twist describes how the local frame rotates around the center-line, being positive for right-handed rotations, negative for left-handed rotations and zero for no rotations (the orange and pink lines mark $\pm\mathbf{d}_1(s,t)$). (a) and (b) are taken from (8).

#### 5. Scaling analysis of growth models

We here investigate our model for the dynamics of growing organs that have similar properties to *Arabidopsis thaliana* roots as described in the methods section. We begin by focusing on the dynamics in the absence of mechanical interactions with the environment. We distinguish between apical sensing, which is theoretically tractable, and local sensing which we analyse numerically. In both cases we estimate the gravitropic turning time $T_0$ and the maximal curvature. In addition, we investigate the scaling of waving patterns on inclined planes for both apical sensing and in the high proprioception regime.

**A. Apical sensing.** The dynamics for roots that have a finite sub-apical growth zone and apical gravisensing are modeled in (7). There, due to the apical sensing the curvature in the growth zone is uniform and obeys the dynamics presented in Eq. S35. When the angle between the tangent at the apex and the direction of gravity is small the dynamics can be approximated by:

$$y'' + (\eta+\gamma)y' + \eta y = 0 \qquad \text{[S36]}$$

which is an ODE of a damped harmonic oscillator, where $y = L_{gz}\kappa^0$, $y' = \frac{dy}{dx}$, $x = \dot{\varepsilon}_g t$, $\eta = \beta L_{gz}/R$ and $\gamma$ is the proprioceptive gain. Damped oscillations then occur for weak damping where $2\sqrt{\eta} - \eta > \gamma$, which occurs within the parameter space explored here (e.g., for $\beta = 0.1$ and $L_{gz}/R = 10$ we have $\eta = 1$, giving damped oscillations for any $\gamma < 1$). In the case of damped oscillations the solution for $y(x)$ is:

$$y(x) = -\frac{\eta \sin(\theta_0 - \theta_g)}{\sqrt{\eta - \left(\frac{\eta+\gamma}{2}\right)^2}} e^{-\frac{(\eta+\gamma)x}{2}} \sin\left(\sqrt{\eta - \left(\frac{\eta+\gamma}{2}\right)^2} x\right) \tag{S37}$$

where $\theta_0$ is the initial angle of the tangent of a straight organ and $\theta_g$ is the angle indicating the direction of the gravitational stimulus. This solution can be translated to the following form of the tip angle over time:

$$\theta^{\text{tip}}(t) - \theta_0 = -\sin(\theta_0 - \theta_g)\left(1 + e^{-\frac{\eta+\gamma}{2}\dot{\varepsilon}_g t}\left(\frac{(\eta - \gamma)\sin\left(\sqrt{\eta - \left(\frac{\eta+\gamma}{2}\right)^2}\dot{\varepsilon}_g t\right)}{\sqrt{4\eta - (\eta+\gamma)^2}} - \cos\left(\sqrt{\eta - \left(\frac{\eta+\gamma}{2}\right)^2}\dot{\varepsilon}_g t\right)\right)\right) \tag{S38}$$

We then define $T_0$ as the time at which $\theta^{\text{tip}}(t)$ reaches to its first maximum:

$$T_0 = \frac{1}{\dot{\varepsilon}_g \sqrt{\eta - \left(\frac{\eta+\gamma}{2}\right)^2}} \arctan\left(\frac{\sqrt{4\eta - (\eta+\gamma)^2}}{\eta + \gamma - 2}\right) \tag{S39}$$

We compare this result to the full non-linear dynamics presented in Eq. S35. As shown in Fig. S3B, we find that the time-scale in Eq. S39 agrees well with numerical integration of the dynamics even for large initial angles. Moreover, for initial angles $\theta_0 \approx \pi/2$ we approximate the maximal curvature using $\kappa^0_{\max} \approx T_0 \cdot \max\{\dot{\kappa}^0\}/3 = T_0\dot{\varepsilon}_g\beta/3R$ as shown in Fig. S3C. The resulting scaling of the maximal curvature, wavelength and amplitude of waving patterns simulated with apical sensing appears in Fig. S4.

**B. Local sensing.** In the lack of analytical solutions for the shape of organs that present sub-apical growth with local sensing, we resort to numerical simulations of the dynamics. We use the algorithm presented in (4) and integrate the growth dynamics until the organs elongate by 10 growth zone lengths, while varying the parameters of the growth model between $0.1 \le \beta \le 0.4$, $0 \le \gamma \le 10.0$ and $L_{gz}/R \in \{5, 10, 20\}$. We characterize the dynamics using the tip angle $\theta^{\text{tip}}(t)$ in a similar manner to apical sensing, and find that for $\gamma < 0.5$ the initial dynamics of $\theta^{\text{tip}}(t)$ can be fitted to damped oscillations with an excellent agreement (see examples in Fig. S3D). Moreover, we find that $\theta^{\text{tip}}(t)$ reaches its first maximum around $T_0 \sim \frac{\pi}{2\dot{\varepsilon}_g}\eta^{-3/5}$ (as in Eq. 7 in the main article), and that the maximal curvature scales like $\kappa^0_{\max} \approx T_0 \max\{\dot{\kappa}^0\} = \dot{\varepsilon}_g T_0 \beta/R$ (see Fig. S3E,F). The resulting scaling of the maximal curvature, wavelength and amplitude of waving patterns simulated with local sensing appears in Fig. S4.

**C. High proprioception.** In Fig. S5, we present the wavelength, amplitude and maximal intrinsic curvature of the waving patterns obtained by simulated organs with proprioception coefficients $0.5 \le \gamma \le 5$. We find that the scaling laws described in Eqs. 9-10 in the main text apply for high proprioception with a few alterations. In the case of apical sensing and high propriocpetion the expression for $T_0$ in Eq. S39 isn't valid as the oscillations become over-damped. We therefore estimate the typical turning time $T_0$ for both apical and local gravisensing using $\dot{\varepsilon}_g T_0 = \frac{\pi}{2}(L_{gz}\beta/R)^{-3/5}$ as done for local gravisensing in the low proprioception regime. We find that the maximal curvature of a free organ in this regime scales like:

$$\max \kappa^0_{\text{f}} \approx \frac{\beta}{\gamma R}, \tag{S40}$$

similar to the maximal curvature described in (9). The maximal curvature of the waving pattern then scales roughly like $0.5 \max \kappa^0_{\text{f}}$ (see Fig. S5A). Moreover, Since in Eq. S40 the maximal intrinsic curvature is inversely proportional to $\gamma$, for very high proprioception the threshold for the initial elastic instability (described later in Eq. S52) is never satisfied. For example, in our simulations the elastic instability didn't take place for $\gamma > 5$ even after 3000 growth time-steps.

#### 6. Normal and parallel decomposition of the growth model

Due to the complexity of the dynamics, which includes active growth processes and passive quasi-static interaction with a tilted plane, we do not present a full analytical analysis of the formation of the waving pattern and skewing angles. Nevertheless, we can gain intuition regarding various aspects of the dynamics by assuming the root remains planar on the tilted substrate. To do so, we begin by describing the geometry of a straight root growing on a tilted plane with tilt angle $\alpha$ and skewing angle $\theta$ (see Fig. S6). We place the origin on the base of the root such that the tilted plane is defined as the $xy$ plane and the normal to the plane is $\mathbf{d}_\perp = \hat{\boldsymbol{z}}$. In these coordinates, the tangent of the organ is: $\mathbf{d}_3 = \sin(\theta)\hat{\boldsymbol{x}} - \cos(\theta)\hat{\boldsymbol{y}}$, and the direction of gravity is: $\hat{\boldsymbol{g}} = -\cos(\alpha)\hat{\boldsymbol{y}} - \sin(\alpha)\hat{\boldsymbol{z}}$. Assuming that the center-line of the root is always planar and embedded in the plane $z = R$, we can define a local coordinate frame on the center-line such that one of its unit vectors aligns with the normal to the plane. As

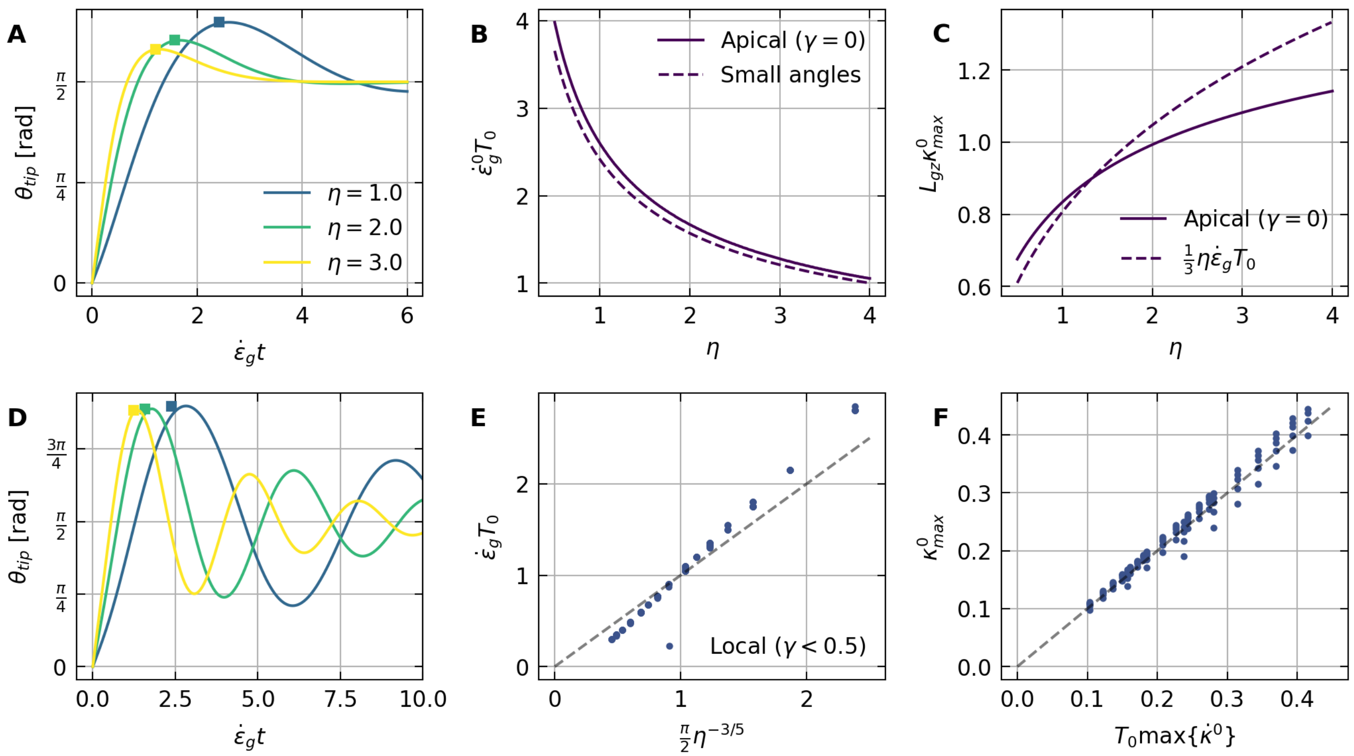

**Fig. S3. Growth dynamic without mechanics:** **A**-**C**: Analysis of apical sensing, based on the analytical results presented in (7). **A**. Example trajectories of $\theta^{\text{tip}}(t)$ for apical sensing with $\gamma = 0$. The squares represent the analytical estimation of $\dot{\varepsilon}_g T_0$ from Eq. S39. **B**. For a large initial angle $\theta_0 \approx \pi/2$ and $\gamma = 0$, we compare between the values for the time $T_0$ describing the first maximum of $\theta^{\text{tip}}(t)$ obtained by both the full non-linear dynamics in Eq. S35 and by the analytical estimation for small angles given by Eq. S39. The values present good agreement despite the large initial angle. **C**. Comparison between the values of maximal curvature obtained by the full non-linear dynamics for $\theta_0 \approx \pi/2$ and $\gamma = 0$ vs. the estimate $\dot{\varepsilon}_g T_0 \beta/3R$, both multiplied by $L_{gz}$. We note that for lower values of initial angles a better approximation is $\dot{\varepsilon}_g T_0 \beta/4R$. **D**-**F**: Analysis of local sensing based on numerical simulations. **D**. Example trajectories of $\theta^{\text{tip}}(t)$ for local sensing with $\gamma = 0$. The squares represent the estimation $\dot{\varepsilon}_g T_0 \approx \frac{\pi}{2}\eta^{-3/5}$. **E**. Numerical estimation of $\dot{\varepsilon}_g T_0$ describing the first maximum of $\theta^{\text{tip}}(t)$ for local sensing when $\gamma < 0.5$ vs. the scaling $\frac{\pi}{2}\eta^{-3/5}$. The dashed line is the identity function added for comparison. **F**. Numerical estimation of $\kappa^0_{\text{max}}$ for local sensing when $\gamma < 0.5$ vs. the scaling $T_0 \max\{\dot{\kappa}^0\} = \dot{\varepsilon}_g T_0 \beta/R$. The dashed line is the identity function added for comparison.

the normal direction is constant all along the organ, this frame is a normal development of the centerline, and we mark the register angle between this coordinate system and the material frame by $\xi$ (Fig. S6) such that:

$$\mathbf{d}_\perp = \sin(\xi)\mathbf{d}_1 + \cos(\xi)\mathbf{d}_2 = \hat{\boldsymbol{z}} \quad \text{[S41]}$$

$$\mathbf{d}_\parallel = \cos(\xi)\mathbf{d}_1 - \sin(\xi)\mathbf{d}_2 = \cos(\theta)\hat{\boldsymbol{x}} + \sin(\theta)\hat{\boldsymbol{y}} \quad \text{[S42]}$$

With a slight abuse of notation, we define the planar component of intrinsic curvature as: $\kappa^0_\parallel = \boldsymbol{\kappa}^{\mathbf{0}} \cdot \mathbf{d}_\perp = \kappa^0_1 \sin(\xi) + \kappa^0_2 \cos(\xi)$ and the normal component of intrinsic curvature as: $\kappa^0_\perp = \boldsymbol{\kappa}^{\mathbf{0}} \cdot \mathbf{d}_\parallel = \kappa^0_1 \cos(\xi) - \kappa^0_2 \sin(\xi)$. These definition are set since according to the right-hand rule it is $\kappa^0_\parallel$ that varies from differential growth parallel to the plane, and it is $\kappa^0_\perp$ that varies from differential growth normal to the plane. If the shape of the organ is planar, $\kappa^0_\perp$ nullifies and the local angle between the tangent and the projected direction of gravity on the plane $\theta(s,t)$ is the integral of the actual parallel curvature:

$$\theta(s,t) - \theta(0,t) = \int_0^s \kappa_\parallel(s',t)ds' \quad \text{[S43]}$$

The skewing angle of a waving pattern is defined to be the average of $\theta$ along the projection of the center-line on the $xy$ plane marked by $\langle\theta\rangle$. We now assume the root is gravitropic and that proprioception is negligible such that $\boldsymbol{\Delta} = \beta\hat{\boldsymbol{g}}$, and focus on the dynamics of the bending components of the curvature, or the projection of the curvature vector on the local cross-section plane: $\boldsymbol{\kappa}^0_{\text{B}} = \kappa^0_1\mathbf{d}_1 + \kappa^0_2\mathbf{d}_2 = \kappa^0_\parallel\mathbf{d}_\perp + \kappa^0_\perp\mathbf{d}_\parallel$. The time derivative of this curvature in the material frame can be written according to Eq. S13:

$$\dot{\boldsymbol{\kappa}}^0_{\text{B}} = \frac{\dot{\varepsilon}_g}{R}(\mathbf{d}_3 \times \beta\hat{\boldsymbol{g}}) = \frac{\dot{\varepsilon}_g}{R}\beta(\sin(\alpha)\cos(\theta)\hat{\boldsymbol{x}} + \sin(\alpha)\sin(\theta)\hat{\boldsymbol{y}} - \cos(\alpha)\sin(\theta)\hat{\boldsymbol{z}}) \quad \text{[S44]}$$

To relate between the dynamics in the frame that rotates with $\mathbf{d}_\perp$ and $\mathbf{d}_\parallel$ to the dynamics in the local material frame from Eq. S44 we must account for the relative rotation between the two frames:

$$\dot{\boldsymbol{\kappa}}^0_{\text{B}}\Big|_{\parallel,\perp} = \dot{\boldsymbol{\kappa}}^0_{\text{B}} + \boldsymbol{\omega} \times \boldsymbol{\kappa}^0_{\text{B}}, \quad \text{[S45]}$$

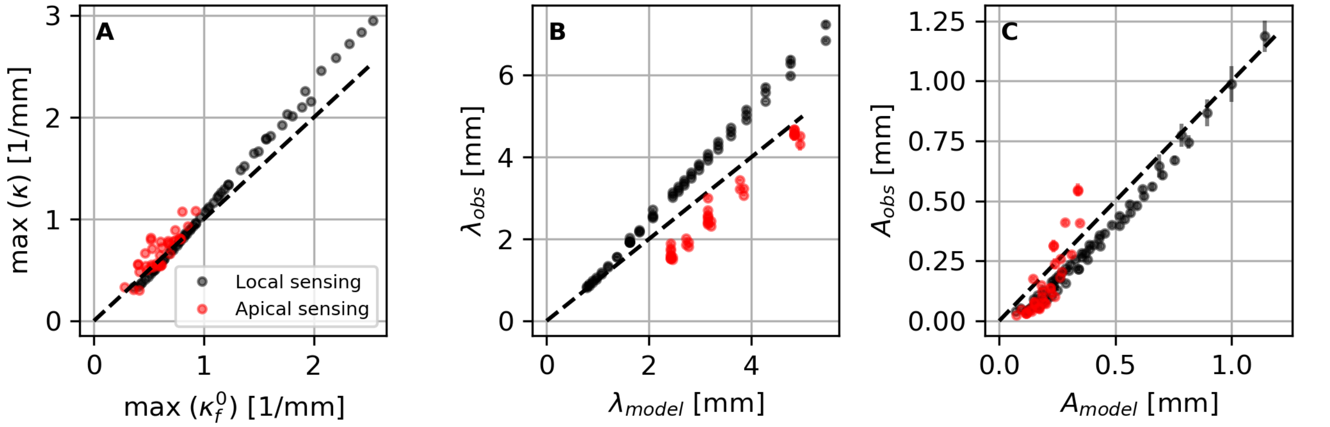

**Fig. S4. Scaling of waving patterns for apical sensing**. Here, we validate our scaling relations (Eqs. 9,10) and compare simulations with apical sensing (with $\gamma < 1$) and local sensing (with $\gamma = 0.1$), each one with its own definition of $T_0$ and the maximal gravitropic curvature of a free organ. **A.** The maximal curvature of the projected shape of simulated waving patterns on the plane $\max(\kappa)$ scales like the predicted maximal curvature of a free organ tilted at the same angle $\alpha$, $\max(\kappa_{\mathrm{f}}^0)$. **B. Wavelength.** Values measured in simulations and experiments $\lambda_{\mathrm{obs}}$ agree with model predictions $\lambda_{\mathrm{model}}$ (Eq. 9), scaling like the gravitropic turning timescale of freely growing organ $T_0$. **C. Amplitude.** Values measured in simulations and experiments $A_{\mathrm{obs}}$ agree with model predictions $A_{\mathrm{model}}$ (Eq. 10), proportional to the time it takes to make one turn in the waving pattern $0.5cT_0 \tan(\alpha)$ (local sensing $c = 1$, apical sensing $c = 1/3$).

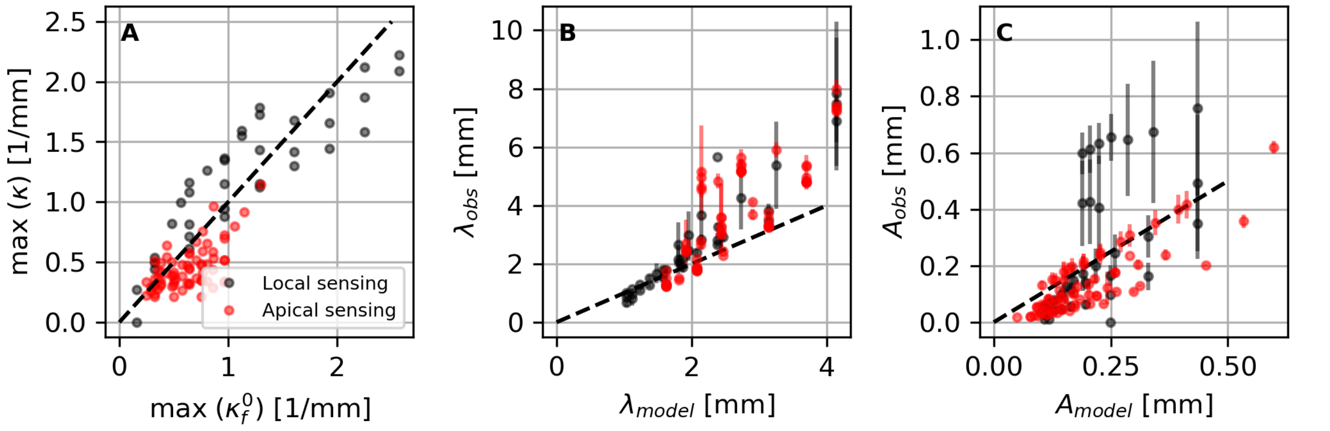

**Fig. S5. Scaling of waving patterns in the high proprioception regime fro local and apical gravisensing** ($0.5 < \gamma < 5$)**.** In this regime the scaling laws presented in Eqs. 9-10 in the main text approximately hold with minor alteration. **A**. The maximal curvature of the projected shape on the plane $\max(\kappa)$ scales like half the predicted maximal curvature of a free organ $\max(\kappa_{\mathrm{f}}^0)$ which now depends on $\gamma$ as described in Eq. S40. **B**. The wavelength of the waving pattern does not depend on the proprioception coefficient $\gamma$ and scales like $2v_g^{\mathrm{tip}}T_0$ as described in Eq. 9. **C**. The amplitude of the waving pattern vs. $0.5cv_g^{\mathrm{tip}}T_0 \tan(\alpha)$ as described in Eq. 10, where now $c = 1/2$ for local sensing and $c = 1/3$ for apical sensing.

where $\boldsymbol{\omega}$ is the relative rotation vector and $\dot{\boldsymbol{\kappa}}_{\mathrm{B}}^0\big|_{\|,\perp} = \dot{\kappa}_{\|}^0 \mathbf{d}_\perp + \dot{\kappa}_\perp^0 \mathbf{d}_\|$. The dynamics of the parallel and normal components of curvature are then given by:

$$\dot{\kappa}_{\|}^0 = \dot{\boldsymbol{\kappa}}_{\mathrm{B}}^0\big|_{\|,\perp} \cdot \mathbf{d}_\perp = \left(\dot{\boldsymbol{\kappa}}_{\mathrm{B}}^0 + \boldsymbol{\omega} \times \boldsymbol{\kappa}_{\mathrm{B}}^0\right) \cdot \mathbf{d}_\perp = \dot{\boldsymbol{\kappa}}_{\mathrm{B}}^0 \cdot \mathbf{d}_\perp + \boldsymbol{\omega} \cdot \left(\boldsymbol{\kappa}_{\mathrm{B}}^0 \times \mathbf{d}_\perp\right) \quad \text{[S46]}$$

$$\dot{\kappa}_\perp^0 = \dot{\boldsymbol{\kappa}}_{\mathrm{B}}^0\big|_{\|,\perp} \cdot \mathbf{d}_\| = \left(\dot{\boldsymbol{\kappa}}_{\mathrm{B}}^0 + \boldsymbol{\omega} \times \boldsymbol{\kappa}_{\mathrm{B}}^0\right) \cdot \mathbf{d}_\| = \dot{\boldsymbol{\kappa}}_{\mathrm{B}}^0 \cdot \mathbf{d}_\| + \boldsymbol{\omega} \cdot \left(\boldsymbol{\kappa}_{\mathrm{B}}^0 \times \mathbf{d}_\|\right) \quad \text{[S47]}$$

where in the last equality we used the vector identity $(\boldsymbol{a} \times \boldsymbol{b}) \cdot \boldsymbol{c} = \boldsymbol{a} \cdot (\boldsymbol{b} \times \boldsymbol{c})$. Since the unit vectors $\mathbf{d}_\perp$ and $\mathbf{d}_\|$ and the curvature vector $\boldsymbol{\kappa}_{\mathrm{B}}^0$ reside in the cross-section plane, only axial rotations such that $\boldsymbol{\omega} \parallel \mathbf{d}_3$ contribute to the second term in the right-hand side of Eqs. S47-S46. These rotations can be expressed by a time dependent register angle $\xi$, and they occur if the organ is twisted elastically or has a time-varying intrinsic twist profile. Marking $\boldsymbol{\omega} = \dot{\xi}\mathbf{d}_3$ and using Eqs. S41,S42,S44,S47,S46 we finally obtain:

$$\dot{\kappa}_{\|}^0 = -\frac{\dot{\varepsilon}_g}{R}\beta \cos(\alpha) \sin(\theta) + \dot{\xi}\kappa_\perp^0 \quad \text{[S48]}$$

$$\dot{\kappa}_\perp^0 = \frac{\dot{\varepsilon}_g}{R}\beta \sin(\alpha) - \dot{\xi}\kappa_{\|}^0 \quad \text{[S49]}$$

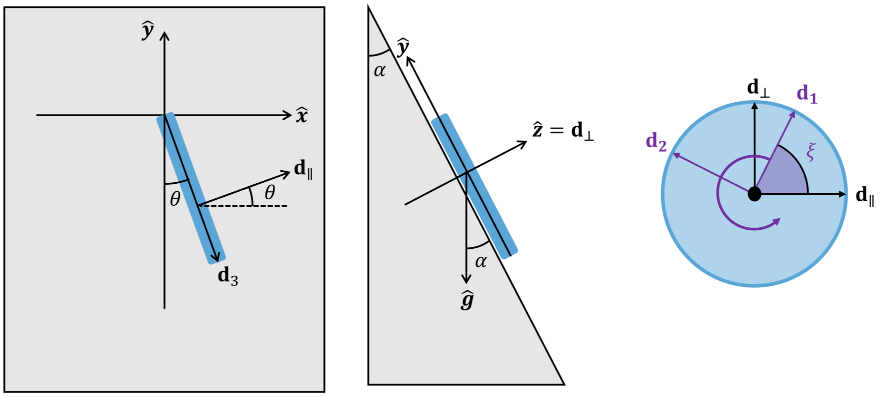

**Fig. S6.** The geometry of a straight root growing on a tilted plane with tilt angle $\alpha$ and skewing angle $\theta$. The local frame $\{\mathbf{d}_{\parallel}, \mathbf{d}_{\perp}, \mathbf{d}_3\}$ on the planar organ, where $\mathbf{d}_{\perp}$ is the normal to the plane, is a normal development of the organ's centerline. We therefore describe the relation between this local frame and the material frame using the register $\xi$ as presented in Fig. 1c.

The first term in the dynamics of the planar component of curvature in Eq. S48 is identical to the dynamics of a free root with an effective gravitropic sensitivity $\beta_{\parallel} = \beta \cos(\alpha)$ and no proprioception, while the second term couples between the parallel curvature and the normal curvature due to twist induced rotations of the curvature vector. The first term in the dynamics of the normal curvature in Eq. S49 then acts as a constant source of curvature with an effective gravitropic sensitivity $\beta_{\perp} = \beta \sin(\alpha)$. If we treat the dynamics of a free root as a damped harmonic oscillation (as shown analytically in the case of apical sensing (7) and numerically for local sensing in SI Appendix 5), these coupled differential equations depict an external drive to the damped oscillations on the plane. Here, a constant source of normal curvature is twisted into the plane due to the elastic interaction with the substrate, which in turn creates a waving or coiling pattern in the mature zone on the plane.

#### 7. Transitions between growth patterns

**A. Transition between straight and waving.** In the lack of intrinsic twist, the initial breaking of symmetry in the development of the waving pattern emerges from an elastic instability that results in elastic twist. This instability takes place only if the elastic energy required for twisting is smaller than the total elastic energy that is gained. At the beginning of the dynamics, the organ grows towards gravity and is pushed back by a normal force from the plane, which keeps the shape of the center-line planar on the plane defined by the direction of gravity and the normal to the plane. In this plane, the organ deforms using 3 degrees of freedom: bending, compression and shear, each with its own elastic energy. Our simulations suggest that the main part of the elastic energy is related to bending deformations. To estimate the maximal bending energy in the growth zone before the breaking of symmetry, we take the actual curvature to be zero (a straight rod) and the intrinsic curvature to be the maximal normal curvature in the low proprioception regime using $\max \kappa_{\mathrm{f}}^0 \sim \dot{\varepsilon}_g T_0 \beta \sin(\alpha)/R$ as described in the maix text and SI Appendix 5. This gives (5):

$$E_{\mathrm{bend}} \approx \frac{1}{2}\int_{L(t)-L_{gz}}^{L(t)} B_{11}(\kappa - \kappa^0)^2 ds \approx \frac{\pi}{8} L_{gz} E R^4 (\max \kappa_{\mathrm{f}}^0)^2, \qquad \text{[S50]}$$

where the bending modulus fits a uniform rod of radius $R$ and Young's modulus $E$ so that $B_{11} = E\pi R^4/4$ (5). The elastic energy due to twist in a rod with no intrinsic twist ($\kappa_3^0 = 0$) can be approximated in a similar manner:

$$E_{\mathrm{twist}} \approx \frac{1}{2}\int_{L(t)-L_{gz}}^{L(t)} B_{33}(\kappa_3 - \kappa_3^0)^2 ds \approx \frac{\pi}{12} L_{gz} E R^4 \kappa_3^2 = \frac{\pi}{12 L_{gz}} E R^4 \Delta\xi^2, \qquad \text{[S51]}$$

where accordingly $B_{33} = E\pi R^4/6$, and in the last equality we defined the register angle in the growth zone due to elastic twist by $\Delta\xi = L_{gz}\kappa_3$. Demanding $E_{\mathrm{bend}} > E_{\mathrm{twist}}$ gives a threshold for the elastic instability that relates the maximal deflection angle of the growth zone $L_{gz} \max \kappa_{\mathrm{f}}^0$ to $\Delta\xi$:

$$L_{gz} \max \kappa_{\mathrm{f}}^0 > \sqrt{\frac{2}{3}} \Delta\xi \qquad \text{[S52]}$$

Substituting $\max \kappa_{\mathrm{f}}^0 \sim \dot{\varepsilon}_g T_0 \beta \sin(\alpha)/R$ and $T_0 \sim \frac{\pi}{2\dot{\varepsilon}_g}(\beta L_{gz}/R)^{-3/5}$ for local sensing gives the threshold in terms of the tilt angle $\alpha$:

$$\sin(\alpha_{\mathrm{s}\to\mathrm{w}}) = \sqrt{\frac{2}{3}} \frac{2\Delta\xi}{\pi} \left(\frac{\beta L_{\mathrm{gz}}}{R}\right)^{-\frac{2}{5}} \qquad \text{[S53]}$$

In this model $\Delta\xi$ acts as a free fitting parameter, and in Fig. 3d we took $\Delta\xi = \pi/8$. In our simulations the elastic instability is more complex than described here as the entire mature zone is free to twist elastically, and the shape of the growth zone prior to the first instability depends on the growth history. Nevertheless, as can be seen in Fig. 3d this model captures the transition well.

**B. Transition between waving and coiling.** To gain intuition regarding the transition between waving and coiling patterns, we approximate the full dynamics using a planar organ as described in SI Appendix 6. Since the reaction force from the substrate is normal to the plane of the organ, the parallel curvature $\kappa_\parallel$ is approximately identical to its intrinsic curvature $\kappa_\parallel^0$ (as can be found by projecting Eq. 2 along with our constitutive model in Eq. 25 on the normal direction). After the first breaking of symmetry, the shape of the growth zone is roughly planar with the same curvature that has been accumulated by the normal differential growth $\max \kappa_{\mathrm{f}}^0$ (as described by Eq. 6). As growth continues, differential growth parallel to the plane attempts to realign the organ towards gravity. However, the turn isn't immediate, and the growing organ keeps drifting uphill based on its existing curvature. These competing processes can be seen by writing the dynamics of the apical angle $\theta^{\mathrm{tip}}(t) \equiv \theta(s = L(t), t)$, as illustrated in Fig. 2a and Fig. S6. Following Eq. S43, in the planar model this angle satisfies:

$$\theta^{\mathrm{tip}}(t) - \theta_0 = \int_0^{L(t)} \kappa_\parallel(s,t) ds \qquad \text{[S54]}$$

where we assumed the base is clamped at an angle $\theta_0$. Taking a time derivative of Eq. S54 and using Eq. S48 with no twist, we obtain two competing terms:

$$\frac{d\theta^{\mathrm{tip}}}{dt} = \frac{dL}{dt}\kappa_\parallel^{\mathrm{tip}} + \int_0^{L(t)} \frac{d\kappa_\parallel}{dt} ds = \dot{\varepsilon}_g L_{gz} \kappa_\parallel^{\mathrm{tip}} - \frac{\dot{\varepsilon}_g}{R}\beta\cos(\alpha) \int_{L(t)-L_{gz}}^{L(t)} \sin(\theta(s,t)) ds \qquad \text{[S55]}$$

where we denote $\kappa_\parallel^{\mathrm{tip}} = \kappa_\parallel(L(t), t)$. The first term on the right hand side of Eq. S55 represents the passive orientation drift due to growth of a curved organ (9), while the second term is the differential growth parallel to the plane that has an effective gravitropic sensitivity $\beta\cos(\alpha)$. Coiling occurs when the angular velocity of the root's apex remains positive throughout the dynamics ($d\theta^{\mathrm{tip}}/dt > 0$), whereas waving occurs when this angular velocity becomes negative and the organ manages to reorient towards the projected direction of gravity on the plane. We express the competition between these two processes using their typical timescales: the turning time in a waving pattern can be expresses using the amplitude scaling in Eq. 10 as $T_0 \tan(\alpha)$, and from Eq. S55 we can estimate the typical time-scale for the passive orientation drift as $T_{\mathrm{dr}} = 1/\dot{\varepsilon}_g L_{gz} \kappa_\parallel^{\mathrm{tip}} \approx 1/\dot{\varepsilon}_g L_{gz} \max \kappa_{\mathrm{f}}^0$. The transition between waving and coiling occurs when $T_{\mathrm{drift}} \approx C T_0 \tan(\alpha)$, which gives an equation for the transition angle $\alpha_{\mathrm{w}\to\mathrm{c}}$ with a fitting parameter $C$:

$$\frac{\sin^2(\alpha_{\mathrm{w}\to\mathrm{c}})}{\cos(\alpha_{\mathrm{w}\to\mathrm{c}})} = \frac{4}{C\pi^2}\left(\frac{\beta L_{\mathrm{gz}}}{R}\right)^{\frac{1}{5}} \qquad \text{[S56]}$$

In Fig. 3d we show that this model captures the transition between waving and coiling patterns well with $C = 0.5$.

**C. Additional configuration spaces.** In Fig. S7 we present four configuration spaces for apical and local sensing and for low and high proprioception. The background color in the low proprioception spaces represents the pattern transitions predicted by our models with critical angles $\alpha_{\mathrm{s}\to\mathrm{w}}$ and $\alpha_{\mathrm{w}\to\mathrm{c}}$ (Eqs. S53,S56) in good agreement with simulations. For apical sensing we repeated the calculations according to the estimations of $T_0$ and the maximal curvature shown in SI Appendix 5.A. One can see that as the proprioception gain $\gamma$ rises, the transition angles between straight, waving and coiling roots increase.

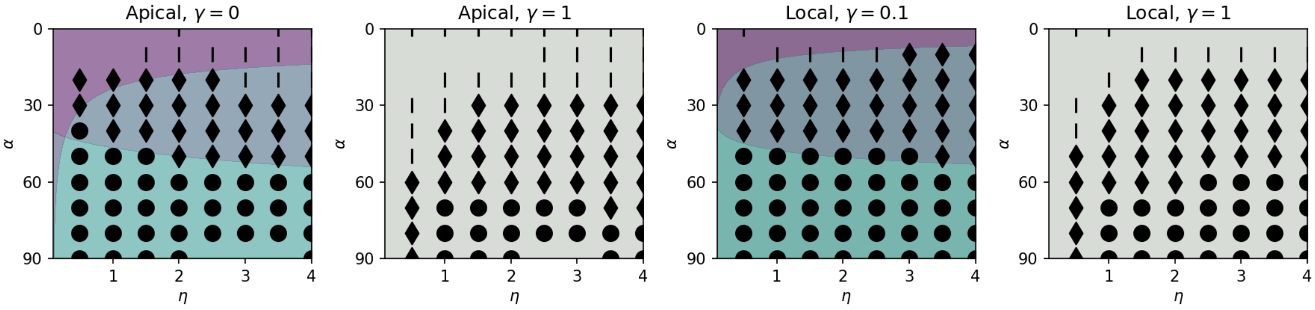

**Fig. S7. Configuration spaces** describing the shape of simulated organs for apical and local sensing in the low and high proprioception regimes. Each Configuration space is presented with respect to varying values of the gravitropic gain multiplied by the slenderness ratio $\eta = L_{gz}\beta/R$ and the tilt angle $\alpha$. Points represent the final configuration in simulations; straight, waving and coiling are represented by bars, diamonds and circles respectively. The background color in the low proprioception spaces represents the pattern transitions predicted by our models with critical angles $\alpha_{\mathrm{s}\to\mathrm{w}}$ and $\alpha_{\mathrm{w}\to\mathrm{c}}$ (Eqs. S53,S56) in good agreement with simulations.

#### 8. Circumnutations

Circumnutations are oscillatory motions of shoots and roots which are linked to differential growth with a direction that rotates around the centerline (3, 10). As described in the methods section (Eq. 19) we model this active process by an additive term in the differential growth vector (3, 4):

$$\mathbf{\Delta}_{CN} = \lambda\left(\cos(\Omega t)\hat{\mathbf{m}}_1 + \sin(\Omega t)\hat{\mathbf{m}}_2\right) \quad \text{[S57]}$$

where $\lambda$ is the circumnutation sensitivity and $\Omega$ is its temporal frequency. Substituting this model into the twits-less curvature dynamics in Eq. S13 gives:

$$\dot{\boldsymbol{\kappa}}^0 = \frac{\dot{\varepsilon}_g}{R}\lambda\left(-\sin(\Omega t)\hat{\mathbf{m}}_1 + \cos(\Omega t)\hat{\mathbf{m}}_2\right) \quad \text{[S58]}$$

Equation S58 can be simplified by marking:

$$\boldsymbol{\kappa}^0_{\mathrm{CN}}(s,t) = \frac{\dot{\varepsilon}_g\lambda}{\Omega R}\left(\cos(\Omega t)\hat{\mathbf{m}}_1(s,t) + \sin(\Omega t)\hat{\mathbf{m}}_2(s,t)\right) \quad \text{[S59]}$$

which allows to rewrite the dynamics from Eq. S58 as:

$$\dot{\boldsymbol{\kappa}}^0 = \Omega\mathbf{d}_3 \times \boldsymbol{\kappa}^0_{\mathrm{CN}} \quad \text{[S60]}$$

This expression resembles the dynamics of twist induced rotations shown in Eqs. S14,S15, which with respect to the normal development frame can be expressed by $\dot{\boldsymbol{\kappa}}^0 = \dot{\xi}\mathbf{d}_3 \times \boldsymbol{\kappa}^0$. This resemblance can also be seen by using the planar geometry presented in SI Appendix 6 while setting $\dot{\xi} = 0$ and assuming a gravitropic and circumnutating root with $\mathbf{\Delta} = \beta\hat{\boldsymbol{g}} + \mathbf{\Delta}_{CN}$. This differential growth model then gives the following dynamics in the material frame:

$$\dot{\kappa}^0_1 = \frac{\dot{\varepsilon}_g}{R}\beta\sin(\alpha) - \Omega\kappa^0_{CN}\sin(\Omega t) \quad \text{[S61]}$$

$$\dot{\kappa}^0_2 = -\frac{\dot{\varepsilon}_g}{R}\beta\cos(\alpha)\sin(\theta) + \Omega\kappa^0_{CN}\cos(\Omega t) \quad \text{[S62]}$$

These relations are similar in structure to those describing gravitropsim with a time varying intrinsic twist profile in Eqs. S49-S48. However, unlike intrinsic twist that rotates the existing curvature with frequency $\dot{\xi}$, the resulting dynamics for twistless cirumnutations described in Eqs. S61-S62 create active curvature of the order of $\kappa^0_{CN} = \dot{\varepsilon}_g\lambda/(\Omega R)$ that rotates with a frequency $\Omega$. Therefore, for certain values of $\kappa^0_{CN}$ and $\Omega$ that resemble the curvature and period of the waving pattern we can expect that the root will present skewing angles in a similar fashion to an intrinsic twist profile. For simplicity, we take $\beta/R$ as the order of magnitude of the gravitropic curvature (following the analysis in SI Appendix 5).

To investigate the effect of our model for circumnutation on the waving patterns we simulated roots with gravitropism, proprioception and circumnuatations, such that $\mathbf{\Delta} = \beta\hat{\boldsymbol{g}} - \gamma R\kappa\hat{\mathbf{N}} + \mathbf{\Delta}_{CN}$, and varied their related parameters between $\gamma = 0.1$, $0 \leq \beta \leq 0.2$, $0.05 \leq R\kappa^0_{CN} \leq 0.15$ and $50\Delta t \leq 2\pi/\Omega \leq 350\Delta t$. We find that simulations of roots with circumnutation and no gravitropism ($\beta = 0$) do not present waving patterns, however the organs develop skewing angles that sometimes result in coiling regardless of the tilt angle. This result may explain why agravitropic mutants and roots grown on a clinostat create coils (11). Simulations with $0 < \kappa^0_{CN} < \beta/R$ result in a waving patterns with low to very low skewing angles ($|\langle\theta\rangle| < 10^o$, see Fig. S8) along with disordered shapes. Simulations with $\beta/R < \kappa^0_{CN}$ result mainly in disordered shapes. We conclude that the model for circumnutations in Eq. 19 may result in skewing angles and coiling, however it does not lead to waving patterns in the absence of gravitropism.

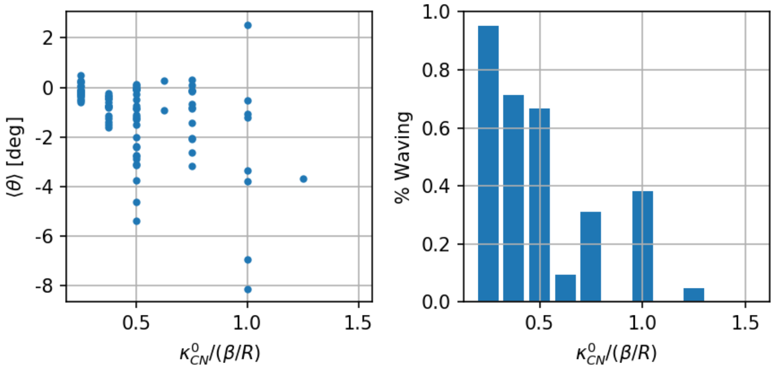

**Fig. S8.** Left: Circumnutations lead to low skewing angles ($|\theta| < 10^o$). Right: The percentage of waving patterns decreases as $\kappa^0_{CN}/(\beta/R)$ rises. The remaining patterns present coiling and disordered shapes.

#### 9. Skewing - additional figures

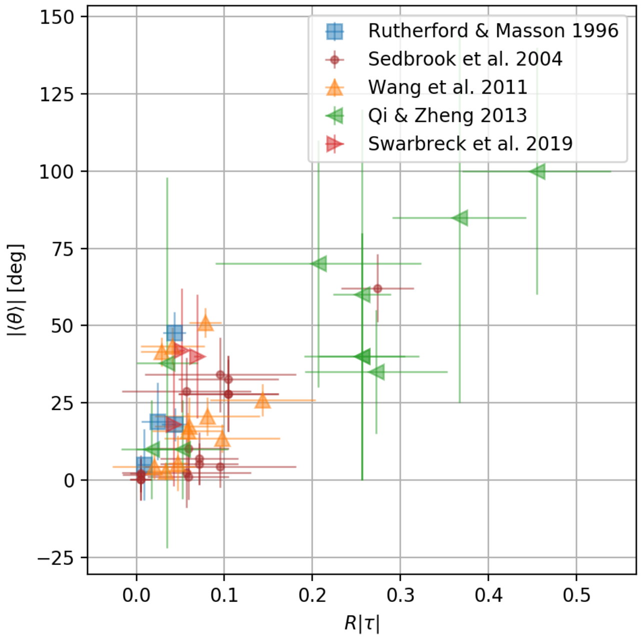

**Fig. S9.** Intrinsic twist vs. Skewing angle - data from experiments, taken from: (12–16).

### Bibliography

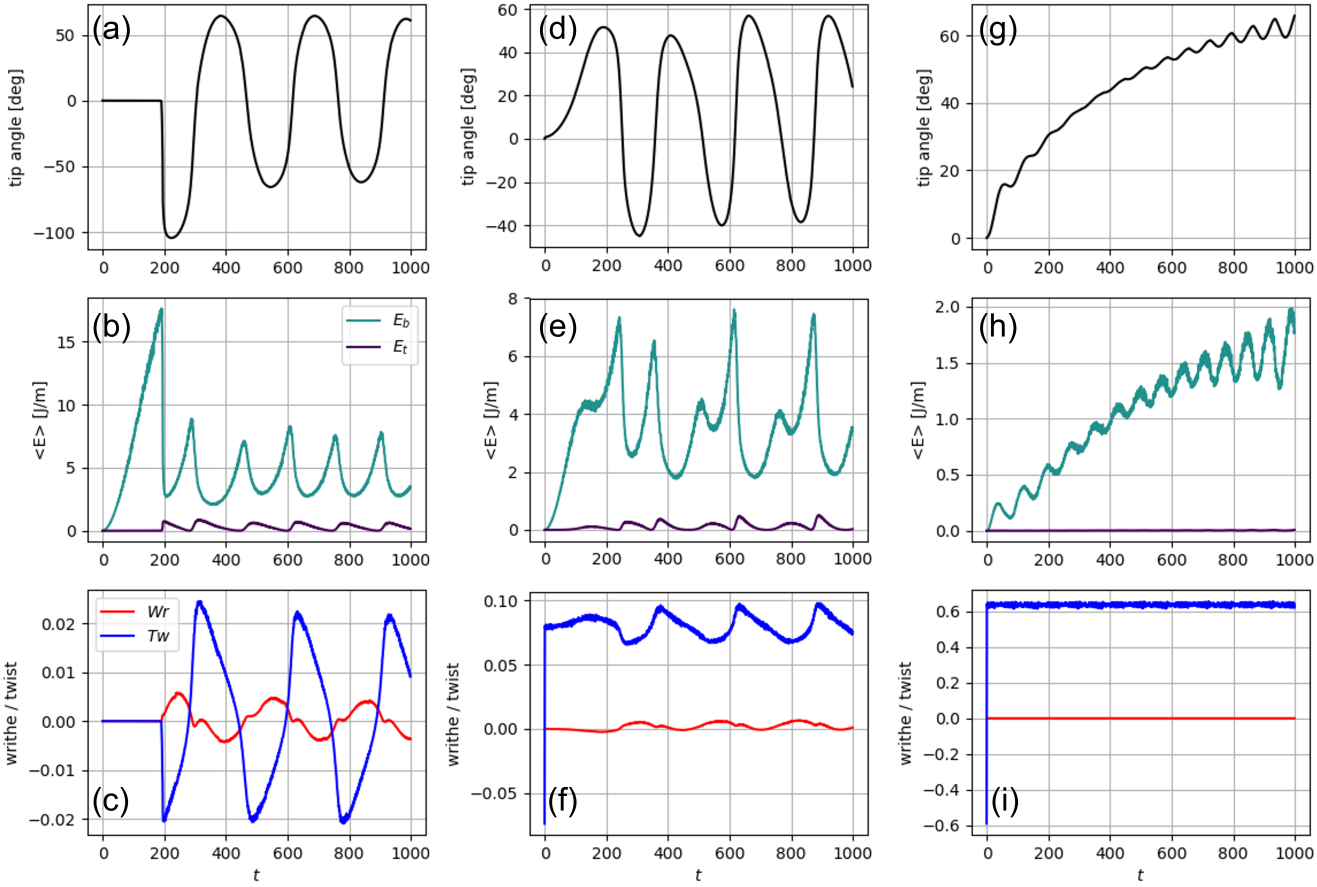

**Fig. S10. Comparison between the development of skewing patterns for various** $\omega T_0$, similar to Fig. 4(d)-(f) for three values of intrinsic twist: $\omega T_0 = 0$ (a)-(c), $\omega T_0 = 1$ (d)-(f), and $\omega T_0 = 8.3$ (h)-(g). Increasing $\omega T_0$ results in larger skewing angles (the average of the tip angle over time) and washes out the waving pattern. Here, $\omega T_0 = 1$ exhibits oscillatory behavior similar to that of regular waving patterns, while $\omega T_0 = 8.3$ transitions to a more monotonic behavior. Note that for a finite value of $\omega T_0$ the initial symmetry is broken by twist without an elastic instability.

16. Stephanie M Swarbreck, Yannick Guerringue, Elsa Matthus, Fiona JC Jamieson, and Julia M Davies. Impairment in karrikin but not strigolactone sensing enhances root skewing in arabidopsis thaliana. *The Plant Journal*, 98(4):607–621, 2019.
